# An *in planta* single-cell screen to accelerate functional genetics

**DOI:** 10.1101/2025.08.06.668818

**Authors:** Tara N. Lowensohn, Will B. Cody, Chun Tsai, Alexander E. Vlahos, Connor C. Call, Xiaojing J. Gao, Elizabeth S. Sattely

**Affiliations:** Department of Chemistry, Stanford University, California 94305; Department of Chemical Engineering, Stanford University, California 94305; Howard Hughes Medical Institute, Stanford University, California 94305

## Abstract

Genetic screens in whole plants are a powerful tool for functional genetics. However, elucidating gene function in highly redundant genetic programs such as signaling pathways remains challenging in both model and non-model plants. Here, we report a single-cell screening platform, PIVOT (Protoplast Isolation after Virus Overexpression *in planTa*), to accelerate identification and functional characterization of plant genes. We used *Nicotiana benthamiana* as a heterologous host to test gene libraries arrayed in a single leaf. Two elements of our system made pooled screens possible *in planta*: (1) we harnessed viral superinfection exclusion to ensure single multiplicity of infection per cell during pooled library delivery, and (2) we engineered a cell surface protein as a phenotypic marker for isolating cells of interest from a heterogeneous population. Using this system, we recovered known and new regulators of cytokinin signaling from an *Arabidopsis* open reading frame library. We anticipate PIVOT will be broadly applicable for high-throughput, single-cell functional genetic screening across the plant kingdom.

## Introduction

Identifying genes that promote plant resilience and plasticity in response to biotic and abiotic stress is essential to ensure future global food security. In this context, genome-wide genetic screens are a crucial tool for gene function elucidation in plants. While whole-plant genetic screens involving a quantifiable phenotype and a population of randomly mutagenized seeds are simple in concept, these screens are notably time, resource, and labor intensive. For example, identification of *fls2*, the gene encoding the immune receptor FLAGELLIN SENSITIVE 2 (FLS2), required manual screening over 80,000 individual *Arabidopsis thaliana* seedlings^1^. Genetic screens in plants can be further complicated by polyploid genomes, self-crossing incompatibility, gene redundancy, or the long life cycles of select plants.

We anticipate that many of the challenges associated with traditional whole-plant screening methods could be overcome if screens were instead carried out using individual cells harboring genetic perturbations. In other organismal systems such as yeast and immortalized human cell lines, cell-based screens have revolutionized gene discovery and are now routine^2^. These screens have been particularly powerful for identifying genes that encode receptors responding to molecular inputs^3^. Similarly, we anticipate cell-based screens could be suitable for many plant functional genetics efforts that do not require whole organisms, such as de-orphaning plant receptors responding to ligands involved in growth and defense^4^.

Analogous single-cell screens have yet to be realized in plants primarily because plant cell culture systems are challenging to transform, passage, and select for a phenotype. Despite these limitations, recent reports of pooled screens in plants reveal the potential for high-throughput functional genetics. For example, libraries of pooled transcriptional enhancer or terminator sequences have been introduced to *Nicotiana benthamiana* leaves, and bulk RNA-sequencing has revealed new genetic elements that boost transcription^5,6^. In more rare cases, pooled libraries have been directly delivered to protoplasts (plant cells devoid of the cell wall). In one example, a cell death-based readout in transformed wheat protoplasts enabled the rapid identification of novel resistance genes and their avirulence gene partners^7^.

Building on this work, we sought to develop a pooled, cell-based, genetic screening platform in plants that could be used for the functional analysis of plant proteins. In contrast to our prior work^8,9^ analyzing enzyme-encoding gene sets in whole leaves without close regard to multiplicity of infection (MOI), we anticipated functional analysis of genes that encode other classes of proteins (e.g., receptors and signaling pathway components) would require assessing gene candidates individually, one per cell. Thus, to realize a cell-based genetic screening platform that moves beyond canonical “one gene query per one plant” screening, two technological advances were necessary: (1) an efficient gene delivery method that results in the expression of libraries of gene candidates with minimal MOI and (2) a cell sorting method to enable single-cell phenotypic analysis.

Here, we describe PIVOT (<u>P</u>rotoplast Isolation after <u>V</u>irus <u>O</u>verexpression *in plan<u>T</u>a*), a first-in-class, pooled, high-throughput *in planta* screening platform to accelerate functional genetics and protein discovery. As proof-of-concept, we tested the ability of PIVOT to identify proteins from *Arabidopsis thaliana* that function in cytokinin signaling. Our approach successfully retrieved both known activators and new cytokinin response factor (CRF) proteins involved in cytokinin signaling. Using PIVOT to recapitulate protein function in the well-characterized cytokinin pathway serves as a model for future discovery of proteins in plant signaling pathways, and we envision this screen could be adapted to survey a broad range of plant protein functions, for example response to small molecules.

## Results

### Designing a platform for screening pooled gene libraries in *Nicotiana benthamiana*

We sought to develop a high-throughput genetic screening platform in plants that would enable assessment of candidate plant proteins involved in perception and transmission of small molecule-mediated signals. Many signaling pathways are widely conserved across plant species, and it has been demonstrated that molecular readouts (e.g., transcriptional reporters) for such pathways are responsive across species^10,11^. Thus, we anticipated a wide range of pathways could be assayed using a single heterologous host. To circumvent plant cell-culture transformation challenges, we leverage existing whole-plant transient expression tools widely used in the model host *Nicotiana benthamiana* and only generate protoplasts at the final screen stage (Figure 1). Our screening strategy involves testing pools of protein-encoding gene candidates in individual cells of an intact leaf, followed by protoplast isolation for cell-level analysis to expedite phenotypic screening.

**Figure 1.**
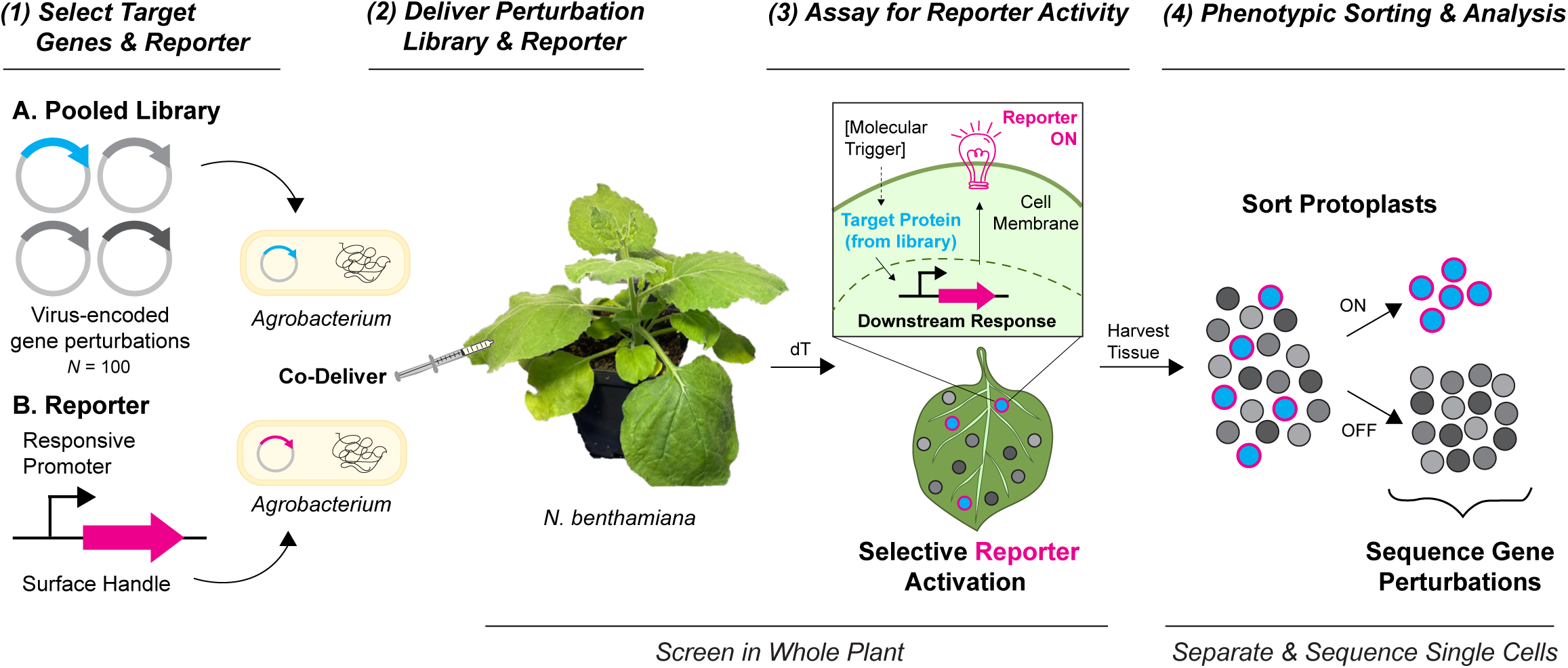
PIVOT workflow for pooled genetic screens in *Nicotiana Benthamiana*. (Steps 1 & 2) *Agrobacterium* strains harboring virus-encoded gene perturbations or a selectively responsive reporter are co-delivered into a mature *N. benthamiana* leaf. (Step 3) Following an incubation period, some gene perturbations cause reporter activation. (Step 4) Single cells (protoplasts) are harvested from leaf tissue, sorted by reporter activity, and sequenced to link phenotype to gene perturbation.

A general overview of our workflow for single-cell genetic screening in *N. benthamiana* is as follows: (1A) Select genes of interest and generate a library of barcoded constructs to perturb (overexpress or knockdown/out) protein-coding gene candidates. (1B) Construct a transcriptional reporter that consists of a cell surface-localized protein expressed under control of a biologically relevant, selectively responsive promoter. (2) Co-deliver the pooled gene library and the transcriptional reporter into *N. benthamiana* leaves. (3) Perform a biological assay in leaves (e.g., incubation, treat with an elicitor, etc.) to assess the role of candidate proteins in modulating transcription of the reporter as a molecular phenotype. (4) Isolate protoplasts, separate based on reporter activity, and sequence cell populations to identify barcoded gene perturbations that alter the reporter activity as a proxy for the phenotype of interest (Figure 1).

### Exploiting viral superinfection exclusion for pooled library delivery

We first turned our attention to developing a gene delivery method that would enable pooled delivery of 10^2^-10^4^ gene candidates to a single *Nicotiana benthamiana* leaf with low MOI per cell (Figure 1, Steps 1 & 2). Importantly, we aimed for each cell to harbor a single gene perturbation to facilitate independent candidate assessment within a pooled screen context. Others recently reported using *Agrobacterium* (*Agro*)-mediated transient expression with canonical binary vectors to simultaneously introduce hundreds of strains in *N. benthamiana* leaves, but to maintain single infections events per cell, the optical density at 600nm (OD_600_) used for *Agro* delivery must be very low (OD_600_ = 0.0025)^12^. An unwanted consequence of this strategy is that a majority (90%)^12^ of the cells in the leaf go untransformed, reducing the experimental output and increasing the amount of infiltrated tissue necessary to ensure robust library coverage.

Alternatively, at high *Agro* OD_600_, we and others have observed that co-infiltration of *Agro* strain libraries results in multiple infections per cell, following a Poisson distribution, where the number of infection events correlates to the *Agro* concentration delivered to a *N. benthamiana* leaf^13^. Here, we qualitatively confirmed this high MOI when using the pEAQ^14^ vector (Figure 2A, top). We used a simple model wherein three *Agro* strains harboring unique fluorescent proteins (FPs) were delivered to a *N. benthamiana* leaf at a final *Agro* OD_600_ of 0.1 (high) or 0.001 (low) (Figure 2B). The Poisson distribution model predicts that the fraction of cells harboring multiple infections increases linearly with the individual strain *Agro* OD_600_; for example, 10% of cells are estimated to be co-infected when two strains are delivered at an *Agro* OD_600_ of 0.01 each, and this figure increases to 50% of cells with individual strain *Agro* OD_600_ of 0.1^13^. For the pooled pEAQ cargo delivered at high OD_600_, we observed co-expression of multiple FPs in single *N. benthamiana* cells (Figure 2C, pEAQ, OD_600_ = 0.1), while at low OD_600_ of *Agro* delivery, we observed sparse single infection events per cell (Figure 2C, pEAQ, OD_600_ = 0.001). These data confirm that single-strain infection events per cell are achievable with pEAQ at low *Agro* OD_600_ delivery, but indeed, at the expense of diminishing the total number of transformed cells.

**Figure 2.**
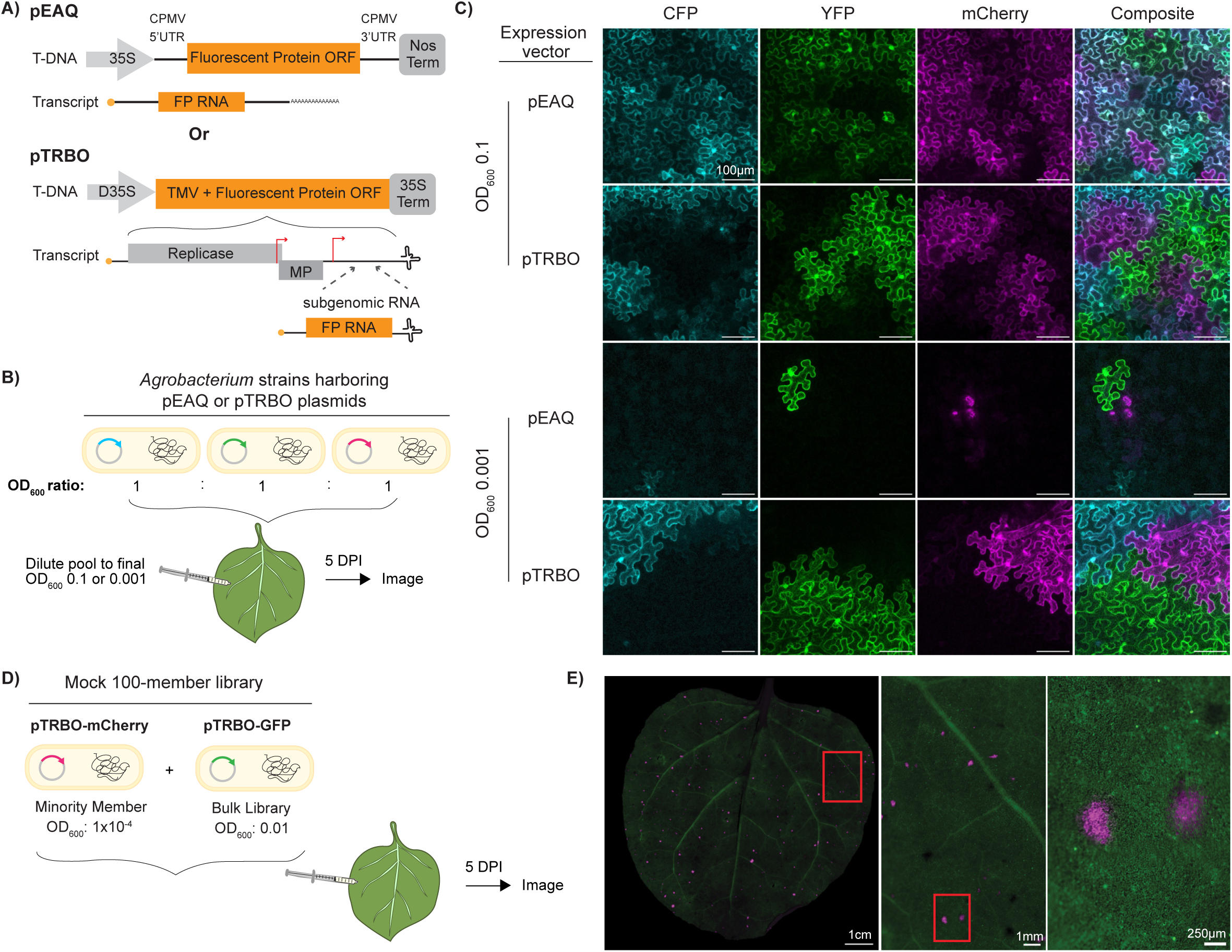
Virus-based open reading frame (ORF) expression enables pooled library delivery *in planta* while limiting the expression of multiple ORFs per cell. (A) Abbreviated plasmid maps showing *Agro*-delivered Transfer DNA (T-DNA) and in-leaf produced transcript from pEAQ and pTRBO vectors. pEAQ T-DNA directly encodes a fluorescent protein (FP) ORF flanked by cowpea mosaic virus (CPMV)-derived untranslated regions (UTRs), while pTRBO T-DNA encodes tobacco mosaic virus (TMV) with a FP ORF inserted after an engineered 3’ subgenomic promoter (red); MP = movement protein. (B) Workflow for pooled FP delivery using either pEAQ or pTRBO vectors: individual FP *Agro* strains were combined at equal 600 nm optical densities (OD_600_), and the mixture was diluted to high (0.1) or low (0.001) final OD_600_ for delivery into *N. benthamiana* leaves. (C) Confocal microscopy of *N. benthamiana* tissue five days post infiltration (DPI) of pooled FP delivery. (D) Workflow for pooled delivery of a mock 100-member library: *Agro* strains harboring pTRBO-mCherry (minority member; OD_600_: 1×10^−4^) and pTRBO-GFP (bulk library; OD_600_: 0.01) were mixed for pooled delivery into *N. benthamiana* tissue. (E) Widefield stereoscope images at increasing magnification of *N. benthamiana* leaf five DPI with mock 100-member FP library.

We wondered if we could instead use a virus that exhibits superinfection exclusion (SE) to achieve single gene perturbations per cell while maximizing infected tissue area. SE is a naturally occurring mechanism enacted by monopartite viruses to prevent secondary, competing infections in the same cell. SE has been observed for bacteria^15^-, plant^16^-, and animal^17^-associated viruses, and notably, has been used to protect crops from viral infection, a practice known as cross-protection^18^. The SE mechanism has been reproduced in experimental settings for many viruses, including tobacco mosaic virus (TMV)^19^, a single-stranded, positive-sense RNA plant virus. A TMV-derived vector, pTRBO (<u>T</u>MV <u>R</u>NA-<u>b</u>ased <u>O</u>verexpression), has been highly optimized for heterologous protein expression in plants and compatibility with *Agro*infiltration^20^. pTRBO contains a viral replicase, a movement protein, and an engineered 3’ sub-genomic promoter to drive gene-of-interest expression (Figure 2A, bottom).

Here, we tested whether pTRBO could be used as a pooled gene delivery tool to achieve both low MOI and high coverage of transformed tissue. Compared to the pEAQ vector, when *Agro* strains harboring FPs encoded in pTRBO vectors are pooled for delivery, single MOI is maintained regardless of the *Agro* delivery OD_600_ because of viral SE (Figure 2C, pTRBO; Supplementary Figure 1). In addition, due to the TMV movement protein that facilitates virus movement between cells, transformation is highly efficient; in this experiment, we were unable to detect any visibly untransformed cells when pTRBO was used, even at a low inoculum concentration (Figure 2C, pTRBO, OD_600_ = 0.001). These data demonstrate *Agro*-launched pTRBO is a suitable vector for pooled gene perturbation delivery that results in single-strain infection at the cellular level while maximizing the transformed cellular space available for downstream phenotypic screening.

We next explored the dynamic range of this virus-based pooled delivery tool to determine how many unique gene candidates introduced as a pool could be detected in a single whole-leaf infiltration. We created a series of mock libraries using defined ratios of two pTRBO-FPs (mCherry and GFP) as proxies for individual gene candidates and used microscopy to visualize FPs as the virus spreads across the leaf (Figure 2D). At five days post-infiltration (DPI), *in planta* expression of two pTRBO-FPs appears to reflect the 1:100 *Agro* OD_600_ ratio (Figure 2E, left). Microscopy analysis confirmed superinfection exclusion was maintained during pooled delivery so that each cell contains a single gene perturbation, while virus movement across the leaf generates pockets of replicates for each gene perturbation (Figure 2E, right; Supplementary Figure 2B).

When this model library was repeated with higher FP ratios, the minority member remained detectable visually in delivered ratios of up to 1 in 10,000 (Supplementary Figure 2C) and quantitatively by barcode sequencing in mock libraries simulating up to 10^6^ members (Extended Data Figure 1). Thus, we anticipate thousands of unique gene queries can be simultaneously delivered but individually assessed in a single *N. benthamiana* leaf (Supplementary Note 1). Taken together, these results highlight SE-enacting, viral genome-encoding vectors like pTRBO are suitable platforms for pooled delivery of up to 10^5^ candidates per leaf with low MOI.

### Candidate representation *in planta* reflects pooled virus library with ORFs of similar size

Qualitative microscopy analysis of leaves indicated single MOI and high transformation coverage using pTRBO. We next wanted to quantify library member representation via barcode sequencing. We generated a library of barcoded pTRBO-CFP constructs for pooled *Agro* transformation and delivery to an *N. benthamiana* leaf (Figure 3A). At five DPI, CFP expression maximized the area of infected leaf tissue (Figure 3B).

**Figure 3.**
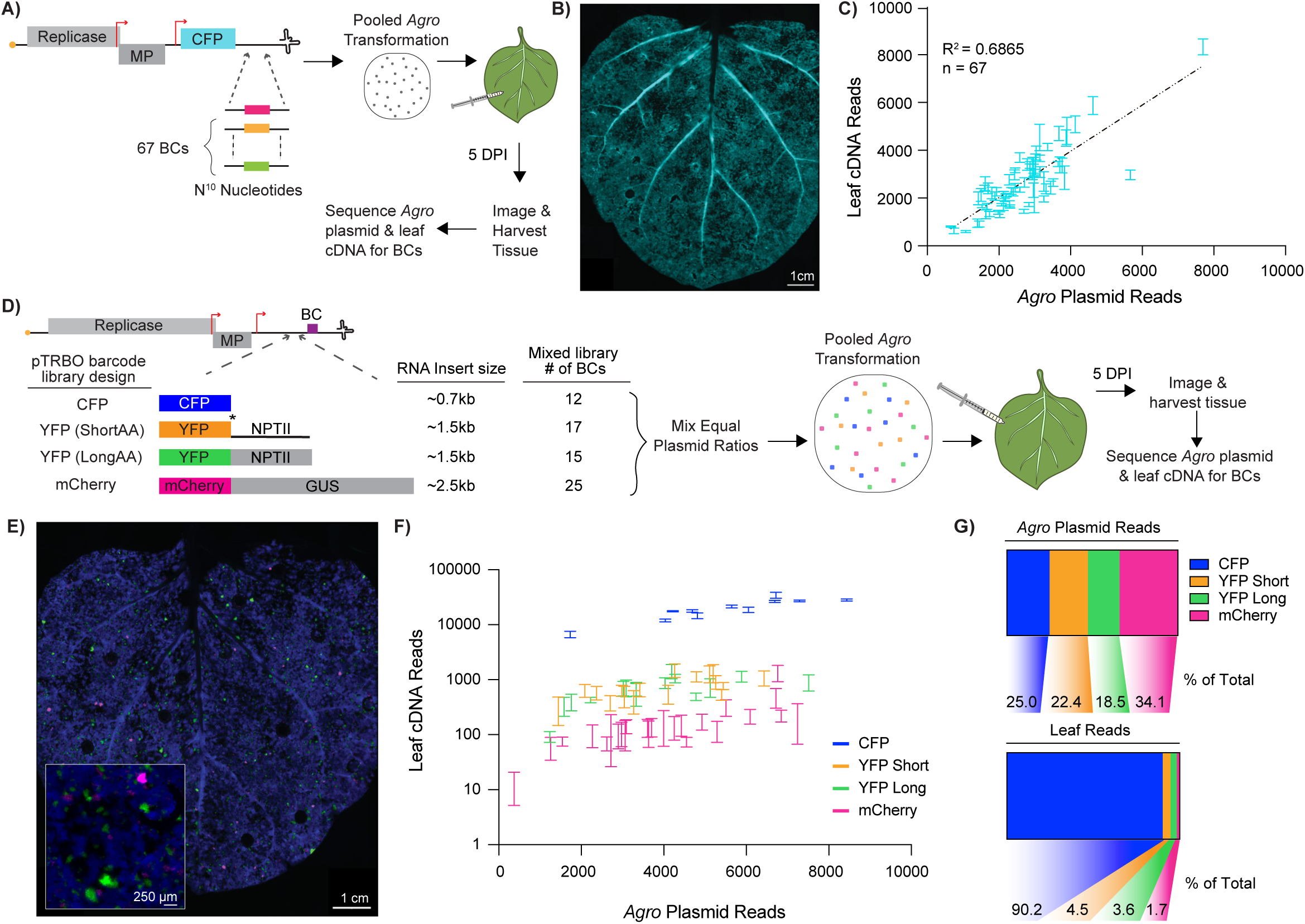
Quantification of representation dynamics for virus-delivered ORF library members. (A) Workflow for quantification of in-leaf barcodes for viral-delivered, equal ORF length library: 67-individually barcoded pTRBO-CFP plasmids were pooled for transformation into *Agro* and infiltrated into *N. benthamiana* (OD_600_ = 0.01). Leaf RNA was extracted five DPI. Plasmid DNA and leaf-derived cDNA were sequenced for barcode quantification. (B) Widefield stereoscope image of pTRBO-CFP-infiltrated *N. benthamiana* leaf at five DPI. (C) Plot of barcode reads before infiltration (*Agro* Plasmid) and five DPI (Leaf cDNA). Each mark represents the standard deviation of 3 replicate reads for an individual pTRBO-CFP barcode, n = 67. Dotted line indicates linear regression, R^2^ = 0.6865. (D) Workflow for quantification of in-leaf barcodes for viral-delivered, mixed size library: pools of barcoded fluorescent proteins (FPs) appended with additional RNA cargo were cloned into the pTRBO-vector. An equal plasmid ratio of each pTRBO-FP sub-library was combined for pooled *Agro* transformation and subsequent delivery into *N. benthamiana* (total OD_600_ = 0.01). (E) Widefield stereoscope, merged channel image of *N. benthamiana* leaf five DPI with mixed size pTRBO-FP pooled library. Increased magnification insert in bottom left corner. (F) Plot comparing barcode representation before infiltration (*Agro* Plasmid Reads) and five DPI (Leaf cDNA Reads). Each mark represents the standard deviation of 3 replicate leaf reads for an individual pTRBO-FP barcode. See (D) for sample size of each pTRBO-FP library. (G) Part-of-whole charts demonstrating input (*Agro*, top) and output (Leaf, bottom) representation of each pTRBO-FP sub-library.

To assess how members of the delivered and *in planta* CFP populations compared, plasmid harvested from *Agro* pool (pre-infiltration), and the cDNA generated from infiltrated tissue were both sequenced to quantify unique post-CFP barcodes. *Agro* plasmid reads and infiltrated leaf-derived cDNA reads revealed the 67 individually barcoded CFPs comprising the library were positively correlated and scaled linearly (Figure 3C). These data indicate that an *Agro*-launched pTRBO vector can deliver a gene overexpression library to *N. benthamiana* with a predictable input-to-output (plasmid-to-*in planta*) candidate representation for equal-sized cargo.

However, a multi-member gene overexpression screen likely contains ORFs of varying size, and we surmised ORF size may affect the dynamics of virus-based delivery^21,22^. Indeed, when we assessed barcoded pTRBO constructs harboring a 2.5 kb insert with mCherry fused to the *E. coli* β-glucuronidase (GUS) ORF, the linear relationship between plasmid and in-leaf barcodes was maintained; however, this larger viral-delivered cargo did not maximize the infiltrated leaf space at five DPI (Supplementary Figure 3). These data suggest that cargo size may influence the degree of viral movement.

To assess the effect of ORF size on candidate representation *in planta* for a virus-delivered library of mixed cargo length, we cloned FPs fused with varying RNA insert size into respective pTRBO vectors: CFP alone (0.7 kb); YFP plus neomycin phosphotransferase II (NPTII) (1.5 kb); and mCherry plus GUS (2.5 kb) (Figure 3D). We cloned an additional pTRBO-YFP-NPTII construct with a stop codon between YFP and NPTII (Figure 3D, “short AA”) to determine if translation size posed an additional burden on pTRBO-based expression *in planta.* Despite approximately equivalent ratios of each pTRBO-FP class in the *Agro* plasmid pool, microscopy analysis at five DPI showed that CFP dominated the *N. benthamiana* leaf space, though pockets of YFP and mCherry expression were still detectable (Figure 3E).

Quantification of barcoded-FP transcripts (Figure 3F) corroborated the microscopy analysis: for a set of varied size cargos, the extent of propagation of the viral clone is proportional to cargo length. Indeed, barcoded CFPs comprised 90% of the in-leaf library reads despite comprising only 25% of the delivered *Agro* plasmid library (Figure 3G). Nonetheless, even the longest members of our library were able to spread and be quantified. Interestingly, the two pTRBO-YFP-NPTII constructs (short and long AA) yielded comparable plasmid and in-leaf reads with respect to one another (Figure 3G), suggesting that additional translation load does not affect library member representation of pTRBO-delivered cargo.

Taken together, these results demonstrate that the in-leaf expression of similar-sized candidates can be predicted by the input library composition, while candidate fitness is affected by cargo length when virus-encoded ORFs of mixed sizes are present in a library. Nevertheless, candidates ranging up to 1.8kb in RNA size can all be sufficiently and reproducibly detected when delivered in a mixed-size pool.

### Engineered surface receptor enables plant single-cell phenotypic sorting

Genetic screening at the single-cell-level requires a method to quickly and efficiently isolate plant single cells (protoplasts) harboring a desired phenotype following gene perturbation (Figure 1, Steps 3 & 4). Many single-cell screens in mammalian systems utilize Fluorescence-activated Cell Sorting (FACS) for facile isolation of cell populations of interest. While protoplasts from some plant species are routinely FACS-sorted, including *Arabidopsis* root^23^, the large size (up to 100 µm in diameter) and fragility of *Nicotiana* protoplasts require specialized FACS configurations^24^. In our experience, *N. benthamiana* protoplasts were prone to lysis during FACS, so we opted to explore alternative methods for protoplast sorting.

We hypothesized transient expression of an engineered receptor protein that localized to the cell membrane could enable magnetic-based pulldown of *N. benthamiana* protoplasts harboring the receptor. We chose the HaloTag protein^25^ as the extracellular receptor handle due to its strong covalent interaction with diverse chloroalkane-linked substrates. Our magnetic-activated protoplast sorting (MAPS) strategy uses a biotinylated HaloTag ligand paired with streptavidin-conjugated magnetic beads to mediate pulldown of HaloTag receptor-harboring protoplasts (Figure 4A).

**Figure 4.**
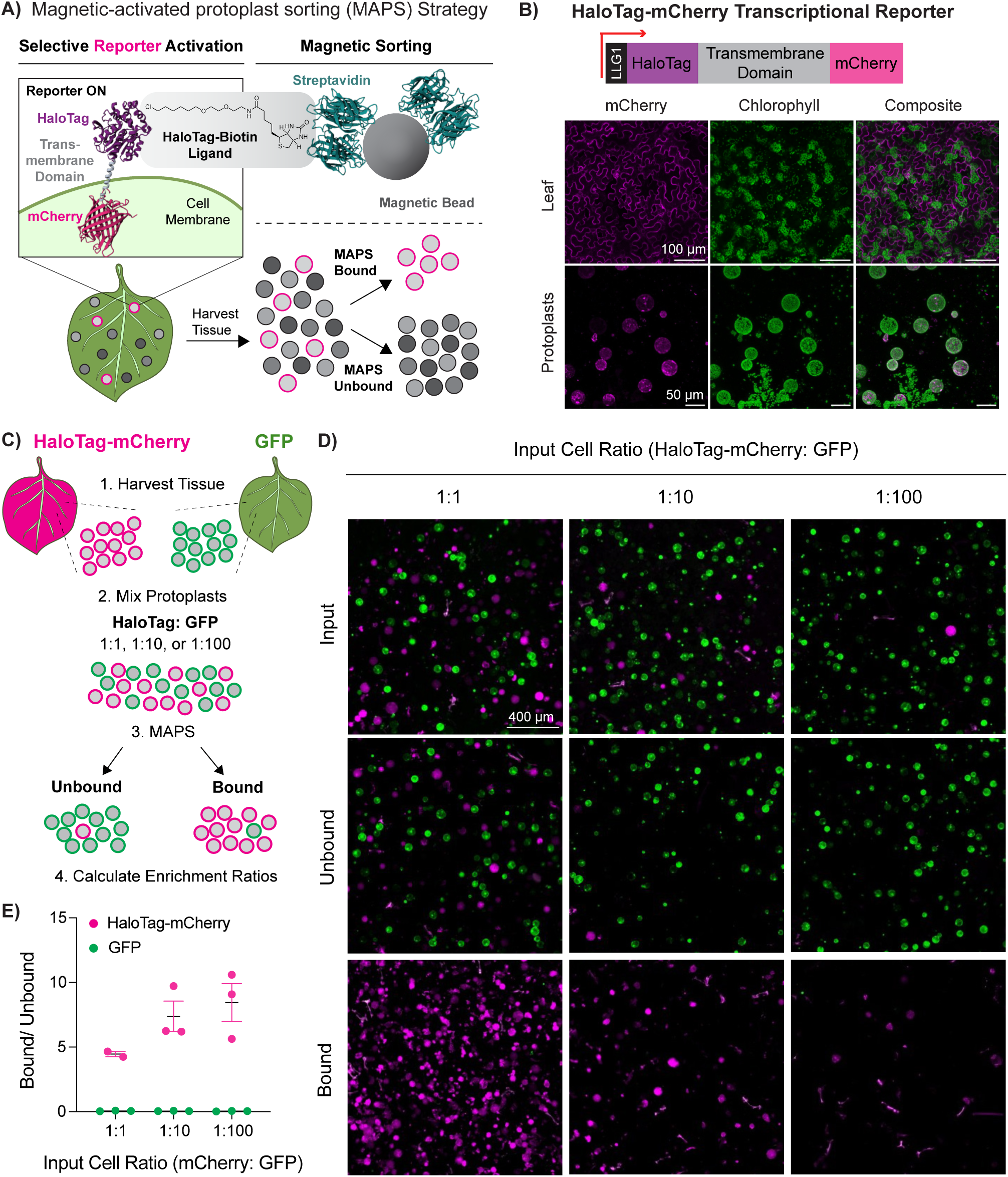
A surface-expressed engineered receptor enables magnetically-activated protoplast sorting (MAPS) of target cells in mixed protoplast populations. (A) MAPS strategy: exogenously added ligand and magnetic beads are used to isolate cells of interest (pink) harboring a selectively expressed cell surface receptor. (B) Abbreviated vector map of a MAPS-capable transcriptional reporter: a promoter of interest (red) drives transcription of HaloTag, which is fused to mCherry for visualization; the signal peptide, LLG1, directs HaloTag to the cell surface, where it is stabilized by a transmembrane domain. Confocal microscope images of infiltrated tissue and protoplasts demonstrating the surface localization of the transcriptionally-encoded HaloTag receptor. (C) Workflow for MAPS selectivity experiments: protoplasts harvested from HaloTag-mCherry (receptor-positive, pink) or GFP (receptor-negative, green)-infiltrated *N. benthamiana* leaves (OD_600_: 0.5) were combined in HaloTag:GFP ratios of 1:1, 1:10, or 1:100, respectively, and then separated using MAPS. (D) Representative fluorescence microscopy images of mixed protoplast populations before (Input) and after (Unbound, Bound) MAPS. See Supplementary Figure 5 for source whole-well images. (E) Enrichment ratios (Bound/Unbound) for GFP versus HaloTag-harboring cells following MAPS. Mean and standard error of the mean enrichment ratios plotted are based on cell counts from 2 or 3 replicates.

We built a MAPS-compatible receptor in which HaloTag is localized to the cell surface by the signal peptide from *Arabidopsis* LORELEI-Like GPI-anchored protein 1 (LLG1)^26^ and anchored in the plasma membrane by a human transmembrane domain (TMD) to stabilize surface expression^27^ (Figure 4B). To assess receptor localization in *N. benthamiana* cells, we *Agro*infiltrated an mCherry-linked version of our HaloTag MAPS receptor under control of the cauliflower mosaic virus (CaMV) 35S constitutive promoter. At four DPI, the HaloTag-mCherry reporter outlines the plasma membrane perimeter of pavement cells, and reporter signal is concentrated at the surface compared to chlorophyll signal distributed throughout the cell (Figure 4B, Leaf). These results indicate the HaloTag reporter is properly trafficked to the plant cell surface in intact tissue. Next, we harvested protoplasts from this surface reporter-infiltrated *N. benthamiana* tissue, and image analysis indicated that a majority (between 94-95%) of recovered protoplasts were indeed transformed and expressing the desired HaloTag-mCherry construct (Supplementary Figure 4). We observed abundant chloroplasts in nearly all transformed cells, indicating our recovered protoplast population is mesophyll-enriched^28,29^ (Figure 4B, Protoplasts).

To determine if MAPS is capable of separating protoplasts based on the presence of a surface-expressed HaloTag receptor, we next generated an artificial mixture of protoplasts with and without the MAPS receptor. CaMV35S-driven HaloTag-mCherry and GFP constructs in *Agrobacterium* were infiltrated into separate *N. benthamiana* leaves; protoplasts with (HaloTag) or without (GFP) the receptor were harvested and mixed at ratios of 1:1, 1:10, or 1:100, respectively, in pools of 10^6^ cells total (Figure 4C).

Figure 4D displays representative images of the mixed protoplast populations before (Input) and after (Bound and Unbound) MAPS. The input images reflect the pre-sort, artificially mixed HaloTag-mCherry:GFP protoplast ratios of 1:1, 1:10, or 1:100, respectively (Figure 4D, Input). Across all input cell ratios, the MAPS bound fraction was dominated by intact HaloTag-mCherry protoplasts, with little to no GFP cells (Figure 4D, Bound). Conversely, GFP cells comprised most of the unbound fraction for the 1:10 and 1:100 ratios (Figure 4D, Unbound). Thus, although HaloTag-mCherry localization is less clear in protoplasts (Figure 4B, Protoplasts), our ability to selectively pulldown reporter-containing protoplasts using MAPS suggests that a substantial portion of the HaloTag protein is present at the cell surface. As an aside, we note that the presence of some mCherry-positive cells within MAPS unbound fraction for the 1:1 ratio might reflect saturation of the magnetic beads at this cell number and ratio.

We quantified the separation capability of MAPS by calculating enrichment ratios for HaloTag-mCherry cells and GFP cells based on cell counts in the bound and unbound fractions. The bound/unbound enrichment ratio for GFP protoplasts was around zero for all input ratios, while the mean enrichment ratio ranged from 4.5 to 8.5 for HaloTag-mCherry cells (Figure 4E). These data demonstrate that MAPS can efficiently separate reporter-positive from reporter-null *N. benthamiana* protoplasts from a mixed population.

Next, to determine if MAPS can discriminate between protoplasts with varied levels reporter expression (beyond binary presence or absence), we performed a similar MAPS separation experiment on a mixed protoplast population harboring the HaloTag-mCherry reporter driven by promoters of increasing strength (Extended Data Figure 2A). We aimed to separate minority groups of interest – low and high-strength promoter cohorts – from a medium-strength promoter majority population (Extended Data Figure 2B); indeed, each promoter cohort yielded a statistically significant MAPS enrichment ratio, which increased with promoter strength: the mean bound/unbound ratios hovered around 0.3 for the low strength promoter; ∼1.0 for medium-strength promoter; and ∼2.5 for the high-strength promoter (Extended Data Figure 2C). HaloTag expression (proxied by mCherry fluorescence) corresponded nicely with expected promoter strength and demonstrated a trend similar to MAPS enrichment ratio (Extended Data Figure 2D).

Collectively, these results demonstrate that pairing a cell surface receptor with magnetic sorting enables *N. benthamiana* protoplast isolation from a mixed population when the target cells comprise between 1-50% of 10^6^ total cells, and that relative levels of reporter expression can be distinguished by MAPS enrichment ratios.

### Proof-of-concept Protoplast Isolation after Virus Overexpression *in planTa* (PIVOT) screen for cytokinin activity

With pooled library delivery (pTRBO) and single-cell isolation (MAPS) methods established, we combined these tools to enable a single-cell, functional genetic screen. For our debut PIVOT screen, we chose to probe cytokinin two-component signal transduction as a test case because of broad pre-existing knowledge of key modulators of this phosphorelay pathway including histidine kinase (HK) receptors, histidine phosphorelay (HP) proteins, and response regulators (RRs)^30^, but also room for functional discovery of cytokinin-related proteins (Figure 5A). Moreover, due to conservation of cytokinin signaling across the plant kingdom, we predicted that homologous proteins could integrate with existing cytokinin signaling circuits to function in a heterologous host.

**Figure 5.**
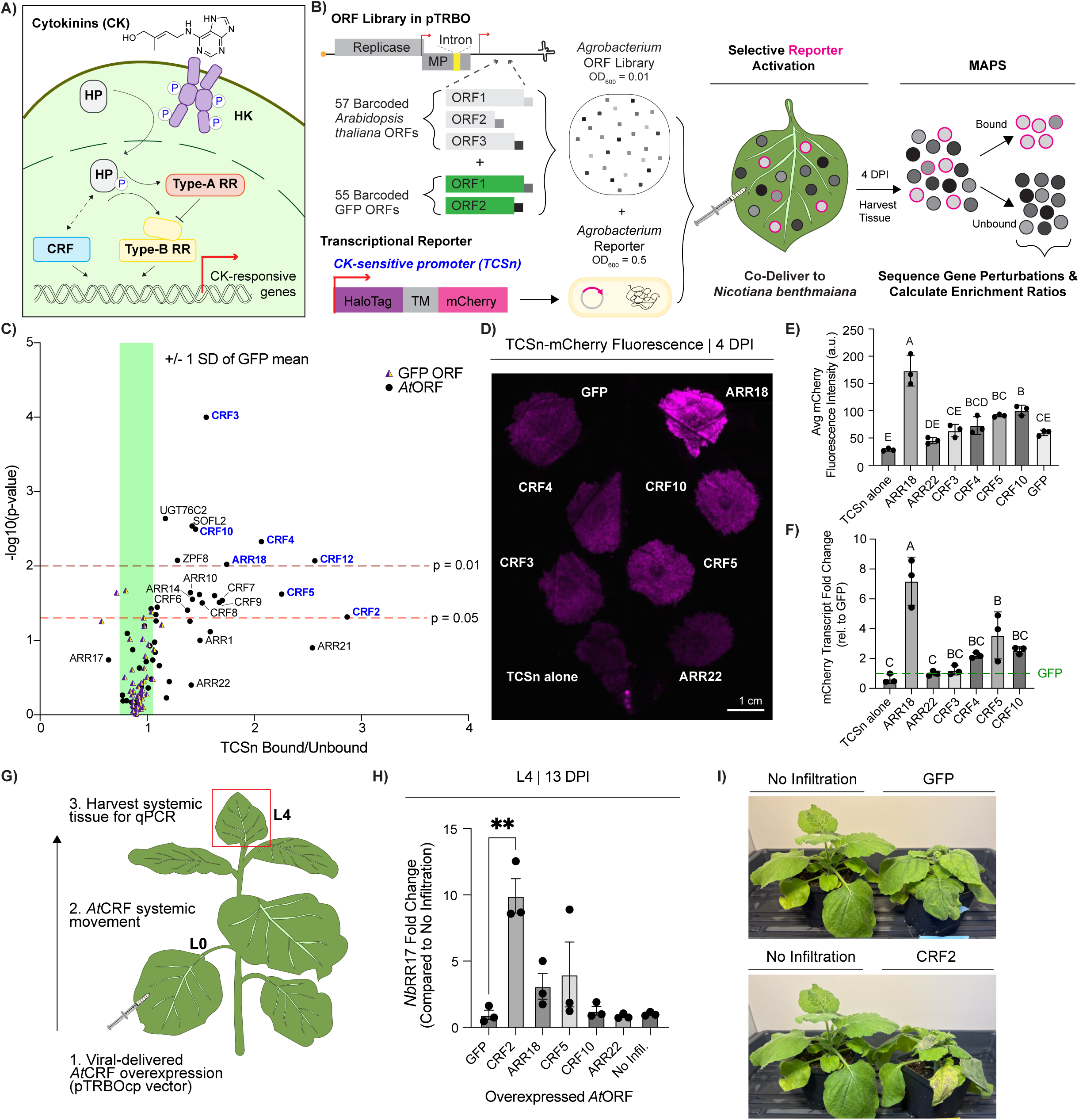
Proof-of-concept PIVOT screen recapitulates CRFs as positive modulators of cytokinin signaling. (A) Abbreviated cytokinin signaling cascade: upon sensing cytokinins (CK), activated histidine kinase (HK) receptors transfer a phosphate group to histidine phosphotransfer (HP) proteins, causing nuclear translocation of HPs. Nuclear HPs can phosphorylate positive (type-B) or negative (type-A) response regulators (RRs). HPs interact with cytokinin response factors (CRFs), but direct phosphorylation has not been shown. CRFs and type-B RRs share some downstream cytokinin-responsive transcriptional targets. (B) Workflow for pooled ORF overexpression screen in *Nicotiana benthamiana*: 57 barcoded *Arabidopsis thaliana* ORFs and 55 barcoded GFP ORFs individually cloned into intron-containing pTRBO vectors were combined for pooled *Agro* transformation and co-delivered into a *N. benthamiana* leaf with a MAPS-capable transcriptional reporter driven by the established cytokinin-sensitive TCSn promoter. Individual *At*ORF activity yields differential reporter activation (pink), generating a marker upon which to sort protoplasts by MAPS. MAPS Bound and Unbound populations are sequenced for barcodes to calculate enrichment ratios for each library member. (C) Volcano plot demonstrating the average of four replicate MAPS enrichment ratios (TCSn Bound/Unbound) for each barcoded GFP (purple triangles, n = 52) or *At*ORF (black circles, n = 52) in the PIVOT screen described in (B). Statistical significance (p-value) derived from independent one-sample, two-tailed *t*-tests comparing the GFP mean MAPS enrichment ratio to that of each barcoded query (See “Calculation of MAPS Enrichment Ratios” in *Methods* for details). Green bar represents ± 1 standard deviation of the GFP mean enrichment ratio (0.8980 ± 0.1553), and the light and dark red lines denote p = 0.05 and p = 0.01 significance, respectively. (D) Representative whole-leaf fluorescent image of post-screen leaf spot assay where *At*ORF candidates were individually assessed for activation of the TCSn reporter; abaxial leaf shown. (E) Mean and standard deviation of three repeat measure fluorescence values derived from the leaf image in panel D. Letters signify statistically significant ANOVA groups from a Tukey’s multiple comparisons test. (F) Quantification of mCherry reporter transcript in tissue harvested from leaf spots shown in (D). Transcript quantifications were normalized to a GFP control. Each dot represents the mean of 3 replicates from an independent experiment. Letters signify statistically significant ANOVA groups from a Tukey’s multiple comparisons test. (G) Workflow for assessing CRF-upregulated genes: ORFs of *At*CRFs were cloned into viral vectors capable of systemic movement (pTRBOcp) and delivered to a *N. benthamiana* leaf. At 13 days post infiltration (DPI) in L0, L4 tissue was harvested for qPCR-based transcript analysis. (H) Mean and standard error of the mean fold change of *N. benthamiana* Response Regulator 17 (*Nb*RR17) for three replicates following systemic overexpression of an *At*ORF. Asterisks indicates significance for two-tailed, unpaired *t*-test between individual query and GFP, ** p < 0.01. (I) Representative images of *N. benthamiana* plants 13 DPI with a virus control (pTRBOcp-GFP) or pTRBOcp-CRF2 compared to non-infiltrated plants.

To carry out the screen, we needed to generate (1) a library of genes to query their role in cytokinin signaling and (2) a mechanism to differentiate genes that activate, repress, or do not mediate cytokinin signaling. We selected ORFs from *Arabidopsis thaliana* genes with known roles directly participating in cytokinin phosphorelay, including type-A and type-B response regulators^30^ (ARRs, A for *Arabidopsis*) (Supplementary Table 1). Further, we included candidates tangentially involved in cytokinin signaling, though not through direct phosphorelay, such as cytokinin response factors (CRFs)^31^ and SOB five-like (SOFL)^32^ genes. We also included circadian rhythm genes and a barcoded GFP library as negative controls, since neither should activate cytokinin signaling. Given that expression of competing pTRBO-delivered cargo is dependent on insert size (Figure 3D-G) we purposefully selected query genes of ORF size within a 1.5kb range to limit bias based on the replication capacity of pTRBO. In total, our pTRBO-ORF library contained ORFs of 57 barcoded *Arabidopsis* gene candidates and 55 barcoded GFPs (Supplementary Table 2). We cloned individual *Arabidopsis* ORFs into an intron^33^-containing pTRBO vector to limit leaky expression in *Agro* without disrupting *in planta* activity of the pTRBO-delivered cargo (Supplementary Figures 6, 7; Supplementary Note 2).

To assess our chosen ORF library for cytokinin-related function, we generated a transcriptional reporter linking the expression of our MAPS cell-surface protein to a previously described cytokinin-sensitive synthetic promoter called TCSn (<u>T</u>wo <u>C</u>omponent signaling <u>S</u>ensor <u>n</u>ew)^10^. We hypothesized that overexpression of known cytokinin signaling activators (such as type-B ARRs) would activate the TCSn promoter, leading to expression of membrane-localized HaloTag (Figure 5B, pink border), whereas overexpression of off-target ORFs would not activate the TCSn promoter (Figure 5B, black border). While the TCSn reporter construct was delivered at high *Agro* OD_600_ (0.5) to ensure as many cells as possible could access inducible reporter expression^13^, the ORF library was delivered at a low *Agro* OD_600_ (0.01) to permit viral spread and produce pockets of individual gene overexpressions (Figure 2E).

Following pooled library and reporter delivery, leaf incubation, and MAPS (Figure 5B), we calculated the enrichment ratio for each ORF based on the number of ORF-specific barcodes in the MAPS bound and unbound fractions. Considering GFP should not biologically contribute to cytokinin signaling, we used the barcoded GFP library as a control group for screen data analysis. A TCSn bound/unbound ratio that is higher than the GFP controls was interpreted to represent TCSn activation and thus extrapolated to indicate ORF-based positive modulation of cytokinin signaling. It is worth noting that using *Agro* to deliver both candidate genes and the reporter construct likely coincides with an induction of cytokinin signaling. We expected *Agro*-derived cytokinin from the *tzs*-harboring GV3101 strain would further contribute to the absolute levels of background TCSn signaling^34^, but that this signal could be enhanced if a pathway protein was transiently overexpressed in a given cell (Supplementary Figure 8).

As predicted, the GFP ORFs generally did not significantly activate the TCSn reporter, and the GFP population maintained a tight distribution around a low TCSn bound/unbound enrichment ratio. The PIVOT screen recapitulated the activity of some known cytokinin signaling regulators. Five of seven tested type-B (positive) ARRs (ARR1, ARR10, ARR14, ARR18, and ARR21) yielded TCSn bound/unbound ratios greater than 1 standard deviation of the GFP mean, though ARR1 and ARR21 were not statistically significant (Figure 5C). Further, the TCSn bound/unbound ratio for six of nine tested Type-A (negative) ARRs (ARR4, ARR6, ARR8, ARR9, ARR16, and ARR17) fell within or below one standard deviation of the GFP mean. TCSn activity for all PIVOT-screened ORFs is reported in Supplementary Table 3.

Interestingly, many cytokinin response factors (CRFs)^31,35^ – AP2 transcription factors harboring an AP2/ERF DNA binding domain and a characteristic CRF protein-binding domain – were amongst the top hits in our screen (Figure 5C). Though CRFs are not known to directly participate in cytokinin phosphorelay, transcriptional changes typically observed following cytokinin treatment are severely altered in *Arabidopsis* CRF mutants, indicating that CRFs are important for regulating cytokinin-responsive genes^31^.

### Post-screen validation highlights CRFs as regulators of cytokinin phosphorelay

To complement the results of our PIVOT screen, we individually validated CRF-mediated TCSn activation. pTRBO-CRFs were co-infiltrated with a TCSn-driven mCherry reporter construct in small leaf spots that were imaged for fluorescence intensity. Initially, we investigated CRFs 3,4,5, and 10 due to their combination of high TCSn bound/unbound ratios and statistical significance compared to the control GFP population (Figure 5C). In *Arabidopsis*, CRF5 was previously known to play a role in cytokinin response^31,36^, while CRFs 3 and 4 have been implicated in embryonic development^37^ and response to cold^38^, respectively. CRF10 controls flowering time in *Brassica napus*^39^ but has not been functionally studied in *Arabidopsis*. We also included ARR18^40^ and ARR22^41^ – known activators and repressors of cytokinin signaling – as positive and negative ORF controls, respectively, in addition to pTRBO-GFP as a control for virus-based delivery.

At four DPI, by visual inspection, mCherry fluorescence was modest for CRFs 3 and 4 and medium strength for CRFs 5 and 10, while the positive control, ARR18, strongly activated the TCSn reporter (Figure 5D). Fluorescence quantification indicated that ARR18, CRF5, and CRF10 are stronger TCSn activators compared to GFP or TCSn reporter-only controls (Figure 5E). As an orthogonal measurement of TCSn activation, RNA harvested from each leaf spot was analyzed for TCSn-driven mCherry transcript levels by quantitative reverse transcription PCR. Fluorescence in the spot assay was corroborated by transcript quantification, with CRF5 and ARR18 yielding significantly more mCherry transcript compared to the reporter-only infiltrated control (Figure 5F).

Though CRF2 and CRF12 were not initially included in validation screening due to borderline statistical significance and low read counts, respectively, precedence from the literature guided us to later follow up on these candidates to understand their behavior in our screen. *Arabidopsis* CRF2 has been shown to play a role in responses to cold, gravity, pathogen, and salt stress, and regulation of reproductive development^35^. Furthermore, CRF2 is the most strongly upregulated CRF in response to cytokinin treatment^36^. On the other hand, the lack of previous knowledge about *Arabidopsis* CRF12 at the time of this study piqued our interest to pursue this candidate further.

In a separate TCSn activation leaf spot assay, CRF12 demonstrated medium-strength mCherry fluorescence, corroborated by a statistically significant increase in mCherry transcript over the pTRBO-GFP control (Extended Data Figure 3A, B). Surprisingly, though CRF2 yielded the highest TCSn bound/unbound ratio in the screen, CRF2 visually and quantitatively failed to demonstrate TCSn activation in this spot assay (Extended Data Figure 3A, B). We noticed CRF2 overexpression in the spot assay resulted in visible cell death at the site of infiltration by four DPI (Extended Data Figure 3C), which could have confounded TCSn activation or our ability to detect mCherry fluorescence. In follow-up spot assays, we assessed TCSn activation at an earlier timepoint (3 DPI) and we reduced the strength of the promoter driving CRF2 expression, which lessened CRF2-induced cell death (Extended Data Figure 3F). The weakest promoter tested (NOS) rescued the expected CRF2-induced activation of the TCSn promoter compared to the GFP control (Extended Data Figure 3D, E), in accordance with PIVOT screen results (Figure 5C). These data highlight the value of cell-based screening for interrogating candidates that could be cytotoxic if overexpressed at a high local concentration or lethal when knocked out (Supplementary Note 3).

### CRF overexpression upregulates type-A response regulator expression

Following validation of CRFs as significant TCSn activators, we explored the role of CRFs in cytokinin signaling outside of the PIVOT screen context. Current models of cytokinin signaling postulate CRFs positively regulate expression of type-A response regulators: type-A ARRs are downregulated in a quadruple *Arabidopsis crf 1,3,5,6* mutant compared to wild-type^42^. Thus, we hypothesized that heterologous overexpression of *At*CRFs in *N. benthamiana* may upregulate expression of endogenous Type-A *Nb*RRs.

We aimed to study the effects of CRF overexpression independent of cytokinin signaling induced by *Agro*infiltration, so we cloned select *Arabidopsis* CRF ORFs into a pTRBO vector containing a viral coat protein (pTRBOcp) that would deliver ORFs to systemic, *Agro*-naïve tissue. Thirteen days after pTRBOcp-CRFs were *Agro*infiltrated into a lower *N. benthamiana* leaf (L0), RNA from systemic tissue (L4) was harvested and analyzed to quantify expression of an endogenous *N. benthamiana* Type-A RR homolog, *Nb*RR17 (Figure 5G). Across multiple systemic leaves in independent experiments, pTRBOcp-CRF5 and pTRBOcp-ARR18 – ORFs that significantly induced TCSn activation in both the PIVOT screen and the leaf spot assay – demonstrated trends suggestive of type-A *Nb*RR upregulation, though not statistically significant (Figure 5H).

Systemic overexpression of *At*CRF2 – which yielded the highest TCSn bound/unbound ratio in the PIVOT screen – demonstrated a 10-fold increase in *Nb*RR17 transcript compared to a GFP control (Figure 5H). CRF2-induced transcriptional alteration led to a striking systemic-wide cell death response (Figure 5I). A similar phenotype was also observed when *At*CRF2 was expressed in local tissue in the leaf spot assay (Extended Data Figure 3C). Identifying the suite of differentially regulated genes in CRF2-overexpressed plants causing this extreme phenotype is an interesting future direction.

In sum, assessing candidates individually for their role in cytokinin signaling supported the data from the screen, providing confidence for utilizing pooled screening approaches for functional genetics. Further, post-screen assays provided experimental evidence for the cytokinin signaling activity of a sequence-predicted, but not yet functionally validated CRF (*At*CRF12), and contributed new evidence supporting the role of CRFs as regulators of type-A RR expression.

## Discussion

Genetic screening in plants has historically required resource-intensive whole-plant methods. In this work, we strove to develop a single-cell screening platform to rapidly increase the pace of plant functional genetics. We harnessed superinfection exclusion from TMV as a delivery mechanism for pooled gene overexpression libraries in plants that enforces single MOI per cell (Figure 2). We demonstrate a single candidate in a pTRBO-encoded mock 10,000-member library can be detected both visually and quantitatively in a single *N. benthamiana* leaf (Supplementary Figure 2C, Extended Data Figure 1A-C). Thus, while our overexpression screen for cytokinin activity included 112 candidates, our data provides evidence that viral vectors could support simultaneous delivery of between 10^4^-10^5^ candidates per leaf (Extended Data Figure 1D-H; Supplementary Note 1).

In our proof-of-concept PIVOT screen, candidate ORFs were individually cloned and validated plasmids were then mixed to create the query library. The approach was done to lessen input library variability while we assessed how leaf-to-leaf variability affected ORF behavior, and our data show that even with biological variation, ORF hits can be statistically differentiated with only 4 replicates (Figure 5C, Supplementary Table 3). For future iterations of a PIVOT screen that are designed to assess larger candidate libraries (10^3^-10^5^ members), it would be desirable to implement bulk cloning methods to easily generate and handle pools of candidate clones. Forthcoming technical advances including enzymatic nucleotide synthesis^43^ and oligo based multiplexed gene assembly methods^44^ could provide welcome solutions to the time, cost, and resource limitations currently plaguing *de novo* gene synthesis. We anticipate combining rapid, facile synthesis methods with viral-based delivery of gene perturbations could provide potential for genome-wide genetic screens to occur in single plants in the future.

When implementing viral delivery in future pooled screening applications, it is important to account for the constraints of these tools. Figure 3D-G demonstrates an inverse relationship between ORF size and in-leaf representation. While this size bias could be avoided in future loss-of-function PIVOT iterations that utilize perturbation strategies that are length-standardized (e.g. hairpin RNAs for gene silencing or guide RNAs for targeted edits), gain-of-function screens involving ORF libraries may require additional optimization. We observed libraries of similar size cargo demonstrate predictable, linear ratios of plasmid-to-leaf barcode representation (Figure 3C, F; Supplemental Figure 3B). These results suggest that ORFs appended with additional RNA cargo to achieve a set length or divided into similar-length sub-pools (e.g. <1 kb, 1-2 kb, etc.) could overcome size-related *in planta* representation bias. Nevertheless, our screen (containing ORFs ranging 1.5 kb in size) identified candidate “hits” both within (ARR18) and nearly two orders of magnitude below (CRF12) expected individual member representation (Supplementary Figure 9).

In addition to establishing virus-based pooled delivery that balances single MOI with high coverage of transformed tissue, we introduce a magnetic-based sorting method for phenotyping at the single plant cell level. Magnetic-activated cell sorting (MACS) has previously been used in plant systems to isolate specific cell types by targeting natively expressed surface markers^45,46^. In a recently disclosed study, MACS was utilized to sort *N. benthamiana* nuclei following transient transformation^12^. Here, we find that cell-surfaced expressed proteins can also be used for magnetic-based sorting of whole protoplasts – which may be too fragile or large for facile FACS sorting – following transient plant gene overexpression.

We integrated pooled gene delivery (pTRBO) and protoplast sorting to enable an *in planta*, pooled genetic screen. As a test case, we screened a 112-member library containing *Arabidopsis thaliana* ORFs for cytokinin activity in the versatile heterologous expression host *N. benthamiana*. Statistical analysis of ORF PIVOT enrichment ratios compared to a control GFP population defined a cohort of putative screen activators for further investigation (Supplementary Figure 10; Supplementary Note 4). Consistent with their respective previously annotated activities^10,47–49^, we found that type-B ARRs 10, 14, and 18 were TCSn activators, while type-A ARRs 4, 6, 8, 9, 16, and 17 did not significantly activate TCSn (Figure 5C, Supplementary Table 3).

Beyond type-B ARRs, cytokinin response factors (CRFs) emerged as consistent activators of the cytokinin-sensitive TCSn promoter during pooled screening (Figure 5C). In addition to cytokinin signaling, CRFs have been implicated in a variety of growth and defense processes^35^. Importantly, assigning individual function to CRFs has been difficult using traditional loss-of-function methods, as CRFs are highly redundant, often requiring triple or quadruple mutants for appreciable phenotype^42^, and some mutant combinations are embryo lethal^31^. Thus, the CRF family represents a case in which PIVOT screening technology could be used to help prioritize candidates whose redundant or deleterious phenotypes can be challenging to resolve using traditional mutagenesis screening.

Our work provides new evidence supporting the role of CRFs in modulating cytokinin signaling-related gene expression. Screen validation experiments determined both CRF5 and CRF12 caused significant TCSn activation (Figure 5D-F; Extended Data Figure 3). Since the TCSn synthetic promoter is derived from type B-RR binding sites, these results corroborate previous findings^31^ that CRFs share downstream gene targets with Type B-RRs (Figure 5A). Notably, *At*CRF12 had not been experimentally analyzed prior to this year when a recent report annotated it as a flowering time regulator^50^. Thus, this work provides the first functional validation for the role of CRF12 in cytokinin response. Furthermore, *At*CRF2 overexpression significantly upregulated RR17, a *N. benthamiana* type-A RR (Figure 5H). These data suggest the conserved role of CRFs in cytokinin phosphorelay regulation across plant families, and that signaling proteins can function in pathways across species.

We anticipate the versatile and modular nature of PIVOT elements (i.e. viral delivery vectors, MAPS-capable reporter, and host choice) could be adapted for diverse plant functional genetics applications. In addition to ORF overexpression, viral vectors can accommodate diverse cargo for eliciting gene perturbations such as guide RNAs^51^, and recently, small CRISPR-like genome editors^52^. These examples present an exciting opportunity to adapt future PIVOT iterations for pooled gene knockdown/knockout screens.

Our proof-of-concept PIVOT screen examined ORF-mediated modulation of a transcriptional reporter, and we envision this workflow will be particularly powerful for assigning gene function in other plant signaling pathways which converge on known gene regulatory elements or where synthetic transcriptional reporters exist. However, we anticipate other phenotypes could be accommodated through clever engineering strategies to regulate MAPS-capable reporters. For example, changes in hormones or marker transcripts following gene perturbation could be captured by linking protein-based small-molecule biosensors^53,54^ or RNA-based sensors^55^ to a MAPS-capable surface handle. Further, the split HaloTag strategy^56^ offers an exciting potential to use MAPS for recording protein-protein interactions. In the future, synthetic genetic^57^ and protein^58^ circuits could permit multiplexed and spatiotemporal control over reporter expression at the transcriptional, translational, or post-translational levels.

A caveat of PIVOT is that our method requires both the reporter and the gene candidates to function properly and integrate with existing signaling pathways in a heterologous host. However, we believe this cross-species screening platform could be amenable for the many signaling pathways that are conserved across the plant kingdom, as demonstrated by our proof-of-concept case investigating cytokinin phosphorelay. Our lab’s previous work elucidating biosynthesis enzymes from across the plant kingdom using *N. benthamiana* as a heterologous expression platform^8,9^ sets a strong precedence for using this host as a screen chassis, and we anticipate PIVOT will permit investigation of ORF libraries from numerous plant species that may not be otherwise genetically tractable.

Nevertheless, PIVOT logic and tools need not be limited to model hosts: dozens of model and crop plants alike have established protocols for protoplast generation, and many characterized, compatible viral vector-host pairs derived from naturally-occurring pathosystems^59^ – including monocot hosts^60^ – could be harnessed for gene delivery. The virus-based pooled gene delivery and protoplast phenotypic sorting techniques comprising the PIVOT platform provide foundational framework for the future realization of genome-wide cell-based screening in plants.

## Acknowledgements

We appreciate feedback we have received from all members of the E.S.S. laboratory (2018-2026). In particular, we would like to thank the following colleagues for their contributions: A. Engel for assistance during the early stages of the project; M. Voges for characterization of the intron used in the pTRBO vector; E. Carlson for assistance with confocal microscopy; D. Wengier and C. McClune for insightful discussions throughout the project; L. Stell (Stanford Medicine) for helpful biostatistics conversations; and C. Liou for comments on the manuscript. This work is supported by the National Science Foundation Graduate Research Fellowship Program (Grant No. DGE-2146755; T.N.L); the U.S. Department of Agriculture National Institute of Food and Agriculture Postdoctoral Fellowship (Grant No. 2022-67012-36700; W.B.C); the Stanford Bio-X Graduate Fellowship (Bowes Fellow; T.N.L.); and the Stanford Synthetic Biology for Sustainability Seed Grant Program.

## Author Contributions

T.N.L, W.B.C., C.T., and E.S.S. contributed to project conceptualization, experimental design, and data analysis. T.N.L, W.B.C., and C.T. performed the experiments. A.E.V, C.C.C and X.J.G. provided the transmembrane domain used to engineer the MAPS-capable receptor; T.N.L. led the manuscript writing with contributions from W.B.C. and E.S.S.

## Data Availability

Raw sequencing data will be made available upon publication.

## Methods

### Cloning Procedures

#### Barcoded fluorescent protein (FP) libraries

Oligo pools containing N10 random nucleotides (barcodes) flanked by primer binding sites were ordered from Integrated DNA Technologies (IDT). The barcodes were PCR-amplified for ∼6 rounds using Invitrogen Platinum SuperFi II PCR Master Mix (ThermoFisher) before insertion into the respective linearized pTRBO-FP backbone using HiFi DNA assembly mix (New England Biolabs). The barcoded pTRBO-FP library was transformed into DH10-beta competent *E. coli* cells (New England Biolabs) and plated on multiple LB-agar plates with 50 ug/mL kanamycin for overnight growth at 37 °C. The next day, between 50-100 colonies across each plate were harvested and combined, and the pooled plasmid DNA was isolated using the Classic Miniprep Kit (Zymo Research). Amplicon sequencing was used to determine the barcodes in the FP library (see *Amplicon Sequencing & Data Processing*).

#### Arabidopsis thaliana Open Reading Frame (ORF) library

ORFs from *At* genes of interest were obtained from The Arabidopsis Information Resource (TAIR). Twist Bioscience synthesized gene fragments with the following syntax for each ORF: [5’ primer binding site]-[ORF]-[adapter sequence]-[10-nt unique barcode]-[3’ primer binding site]. (See Supplementary Table 2 for sequence information.) ORFs were cloned into a linearized, intron-containing pTRBO backbone using HiFi DNA assembly mix (New England Biolabs). Each pTRBO-*At*ORF construct was individually transformed into DH10-beta competent *E. coli* cells (New England Biolabs) and plated on an LB-agar plates with 50 ug/mL kanamycin for overnight growth. A single colony was picked for outgrowth, and the following day, plasmid DNA was isolated using the Classic Miniprep Kit (Zymo Research). Plasmids were sequence-verified with whole-plasmid sequencing (Plasmidsaurus). Equal concentrations of each pTRBO-*At*ORF construct were combined to generate a pooled *At*ORF plasmid library. For the PIVOT screen, plasmid from the pooled *At*ORF library was combined with intron-containing, barcoded pTRBO-GFP library plasmid (for construction, see *Barcoded fluorescent protein (FP) libraries* above) for pooled *Agro* transformation (see *Agrobacterium Transformation and Growth Conditions* below).

#### Systemic-moving AtORF constructs

*At*ORFs were inserted into coat protein-containing viral vectors (pTRBOcp) via restriction cloning. *At*ORFs were PCR-amplified from parent pTRBO-ORF constructs appended with restriction sites compatible with a pTRBOcp vector. *At*ORFs and the pTRBOcp vector were cut with restriction enzymes (New England Biolabs) and ligated with T4 ligase (New England Biolabs), both according to manufacturer’s specifications. *E. coli* transformation, plasmid miniprep, and sequencing procedures were performed in the same manner as described above for the *At*ORF library.

#### Additional sequence information

*(i) MAPS reporter*: the transmembrane domain (see ref. 27, Vlahos et al., 2025) used in our surface reporter construct is as follows (5’-3’): GTGGTATACCCCTGCACGGCCTTGCTGCTGCTCTGCCTCTTCGCCACCATCATCACCTACATCCTC. *(ii) Intron-containing pTRBO vector*: the following intron sequence (see ref. 33, Voges, 2019) was inserted following M198 in the movement protein of the pTRBO vector^20^ (5’-3’): gtaagagtactttatattttactcaccatgtcgtgtcgtaacctcccgtttttttgcgcag.

#### *Agrobacterium* Transformation and Growth Conditions

Individual constructs or pooled plasmids purified from *E. coli* were transformed into *Agrobacterium tumefaciens* (strain GV3101, except for Supplementary Figure 8 – strain LBA4404) using either electroporation (preset *Agrobacterium* protocol on BioRad GenePulser Xcell) or the freeze-thaw method. Transformed cell suspension in medium 523 was shaken at ∼200 rpm at 30 °C for 2 hours, and the post-outgrowth solution was plated on medium 523-agar plates with 50 µg/mL kanamycin and 30 µg/mL gentamicin at 30 °C for 2 days. For individual constructs, a single colony per construct was picked for overnight clonal outgrowth, while for pooled plasmid libraries, 10^2^-10^3^ colonies across multiple plates were harvested and combined to ensure full library coverage. The *GV3101* colonies underwent outgrowth in medium 523 with 50 µg/mL kanamycin and 30 µg/mL gentamicin at 30 °C and 200 rpm for ∼2 days before *Agro*infiltration preparation. Frozen stocks were prepared for each construct by adding glycerol to the culture to a final concentration of 50%.

#### *Agro*infiltration

GV3101 cultures in medium 523 were centrifuged for ∼10 minutes at 4,000xg and resuspended in an infiltration buffer (0.1M MES buffer, pH 5.6, 1M MgCl_2_). Suspensions were vortexed, measured for Optical Density at 600 nm (OD_600_), and delivered to the abaxial side of a 4-5 week-old *Nicotiana benthamiana* leaf using a needleless syringe. Figure captions and experiment-specific methods sections specify OD_600_ values used.

### Plant Growth Conditions

*Nicotiana benthamiana* plants were grown indoors at room temperature for ∼4-5 weeks before infiltration. Briefly, *N. benthamiana* seeds were sown into fertilized soil and covered for two weeks. Two-week-old seedlings were thinned and covered with a humidity dome for an additional week before acclimating to open-air conditions. Following infiltration, plants were incubated in a growth chamber at 22 °C with 50% humidity under a 16/8 day/light cycle until tissue harvesting.

### Confocal Microscopy

*N. benthamiana* tissue samples were prepared and imaged according to previously reported methods (see ref. 13, Carlson et al., 2023).

### Widefield Stereoscope Microscopy for Whole-Leaf Imaging

Fluorescent protein (FP) expression in whole leaves was visualized using Leica THUNDER Imager Model Organism paired with LAS-X software (Leica Microsystems). Clear plastic wrap was used to secure leaves mounted on the microscope stage. Leaves were scanned at 1.25x magnification in “Tilescan” mode in the respective FP channel: CFP (Ex. 436/20 nm; Em. 480/40 nm); YFP (Ex. 510/20 nm; Em. 535/30 nm); mCherry (Ex. 540/40 nm; Em. 593/60 nm); GFP (Ex. 470/40 nm; Em. 525/50 nm). Following mosaic merge, the multichannel image stack was processed using Image J for brightness/contrast adjustments and false coloring.

### Protoplasting of *N. benthamiana* Tissue

Cell wall digestion buffer (DB) was prepared fresh on the day of protoplasting: MES buffer (20mM, pH 5.7), mannitol (400mM), and KCl (20 mM) were combined with cellulase R-10 (1%) and macerozyme R-10 (0.4%) and heated for 10 minutes at 55 °C. The solution was cooled to room temperature before adding CaCl_2_ (10mM) and BSA (1%). The midrib and major veins of the *N. benthamiana* leaf were dissected away and the remaining tissue was sliced into 0.5-1mm strips using a razor blade, transferred into a 6-well plate, and submerged in DB. A desiccator was used to vacuum infiltrate the DB into leaf strips for 30 minutes. Tissue underwent digestion in the dark for 3-4 hours on a plate shaker at 30 °C and 50 rpm. Digested cells were passed through a 100 µm filter, washed in an enzyme-free cell wash buffer (CWB: 20 mM MES buffer, pH 5.7; 20 mM KCl, 10mM CaCl_2_, and 0.5% BSA), and pelleted by centrifugation at 100 x g. A small aliquot of protoplasts resuspended in CWB was stained with Fluorescein Diacetate (FDA) to count live cells using a BioTek Cytation 5 Cell Imaging Multimode Reader (see *Cell Counting and Fluorescence Microscopy of Protoplasts*). Biological replicates were normalized to the desired protoplast concentration (typically between 150-300k cells/mL).

### Magnetic-activated Protoplast Sorting (MAPS)

Immediately following harvesting (see *Protoplasting of N. benthamiana Tissue*), *N. benthamiana* protoplasts were incubated with Promega HaloTag® PEG-Biotin Ligand (2 µL per 2 mL protoplasts) with end-to-end rotation for 1 hour. Protoplasts were then pelletted at 100 x g and washed multiple times with cell wash buffer (CWB) to remove excess ligand. Ligand-bound protoplasts were incubated with CWB-washed, magnetic Dynabeads MyOne T1 Streptavidin beads (ThermoFisher, 20 µL per 100k protoplasts) with end-to-end rotation for 1 hour. An aliquot of bead-bound protoplasts before magnetic sorting was saved as the “Input” fraction. The remainder of the protoplast population was subject to sorting using a StemCell Technologies magnet. Protoplasts bound to the magnet were washed with CWB three times. The first two washes were kept as the “Unbound” fraction, the final wash was discarded, and the remaining protoplasts were resuspended in CWB (“Bound” fraction). Aliquots of “Input”,“Unbound”, and “Bound” populations were imaged using a BioTek Cytation 5 Cell Imaging Multimode Reader when applicable, and the remaining cell pellets were frozen for RNA extraction.

### Cell Counting and Fluorescence Microscopy of Protoplasts

Protoplasts were visualized and counted using a BioTek Cytation 5 Cell Imaging Multimode Reader paired with BioTek Gen5 Imaging software (Agilent Technologies).

#### Protoplast counting

a 50 µL aliquot of protoplasts diluted to 1 mL in cell wash buffer (CWB) was incubated with Fluorescein Diacetate (FDA, ThermoFisher) (2 µL per 1 mL diluted protoplasts) for 5-10 minutes in the dark. Protoplasts were then pelleted by centrifugation at 100 x g and washed with CWB to remove excess FDA stain. Triplicate 200 µL aliquots of FDA-stained protoplasts were loaded in a 96-well black-sided, clear bottom plate (Corning) and imaged using a 4x objective in the GFP channel (Ex. BP 469/35 nm; Em. BP 525/39 nm). Live protoplast counts were gated based on the following post-image processing parameters (Size: 15-100 µm; Circularity: >0.5; Fluorescence: >5000).

#### Protoplast fluorescence microscopy

Following cell sorting, 200 µL aliquots of MAPS Input, Unbound, and Bound fractions were loaded in a 96-well black-sided, clear bottom plate and imaged using a 4x objective in GFP (Ex. BP 469/35 nm; Em. BP 525/39 nm) and RFP (Ex. BP 531/40 nm; Em. BP 593/40 nm) channels. For each fluorescence channel, protoplast counts were gated based on the following post-image processing parameters (Size: 15-100 µm; Circularity: >0.5; Fluorescence: >5000). ImageJ was used for brightness/contrast adjustments and false coloring.

### RNA Extraction and Preparation of complementary DNA (cDNA)

*N. benthamiana* tissue was flash-frozen and ground by hand in 2-mL Eppendorf tubes using plastic pestles. RNA was extracted using a RNeasy Plant Mini Kit (Qiagen) and purified using a RNase-free DNase set (Qiagen). RNA concentration was measured using Nanodrop and normalized across samples (typically to 1000 ng). cDNA was prepared using the SuperScript IV First-Strand Synthesis System (ThermoFisher); random hexamers were used to amplify virus-encoded, barcoded ORF RNAs in library representation experiments (Figure 3, Extended Data Figures 1 and 2, Supplementary Figure 7) and the PIVOT screen (Figure 5C), while oligo dT primers were used to amplify TCSn transcript (Figure 5F) and endogenous *N. benthamiana* RNAs (Figure 5H), respectively, before qPCR analysis.

### Amplicon Sequencing & Data Processing

When applicable, first-strand cDNA was amplified by PCR using library-specific primers (see Supplementary Table 4) with Platinum SuperFi II PCR Master Mix (ThermoFisher) and purified using a DNA Clean & Concentrator kit (Zymo Research). Plasmid or amplified cDNA (25 μL of 20 ng/uL) was sent to Azenta Life Sciences for Amplicon-EZ Next Generation Sequencing. Standard sequencing depth (5 × 10^4^ reads guaranteed) was used, except for the experiment described in Extended Data Figure 1D-F (see also Supplementary Table 5), where custom deep sequencing (on the order of 10^8^ reads) was requested. Paired-end sequencing reads were processed and separated by barcode using Geneious Prime Bioinformatics software. Raw read counts were normalized to between 1-3 × 10^5^ total reads per replicate depending on the experiment before graphical analysis (see figure captions in Supplementary Tables 5-9 for details). A single sample was sequenced for *E. coli* and *Agro* plasmid samples, and figure and table captions indicate the number of sequenced replicates (3 or 4) for leaf cDNA reads.

### Calculation of MAPS Enrichment Ratios

For the proof-of-concept protoplast sorting experiment (Figure 4), MAPS enrichment ratio was calculated by dividing the Cytation 5-derived cell counts (see *Cell Counting and Fluorescence Microscopy of Protoplasts*) for the Bound fraction by the Unbound fraction for each replicate. For the PIVOT screen (Figure 5), Bound and Unbound reads were normalized (see *Amplicon Sequencing & Data Processing*), and for each replicate, the number of Bound reads was divided by Unbound reads. For the PIVOT screen p-values (Figure 5C, Supplementary Tables 3 and 10), a one-sample, two-tailed *t*-test was performed comparing the MAPS enrichment ratios for each *At*ORF and GFP ORF (four replicates) to the mean MAPS enrichment ratio for the GFP population (n = 52, theoretical mean = 0.8980).

### Leaf Spot Assays for Validating PIVOT Screen Hits

*At*ORFs chosen for further inquiry following the PIVOT screen were individually transformed into *Agro* GV3101. Each pTRBO-encoded ORF *Agro* strain was mixed with an *Agro* strain harboring the TCSn-mCherry reporter construct at *Agro* OD_600_ values of 0.1 and 0.5, respectively, and infiltrated in a ∼15mm diameter spot on a 4-6-week-old *N. benthamiana* leaf. At four DPI, the abaxial side was imaged using an Invitrogen iBright FL1500 Imager (ThermoFisher). ImageJ was used for brightness/contrast adjustments, false coloring, and fluorescence quantification. After imaging, three 0.7 cm leaf punches for each construct were harvested for ensuing RNA extraction and qPCR analysis.

### Systemic Overexpression of CRFs in *Nicotiana benthamiana*

Select CRFs and ARR controls were cloned into coat protein-containing virus vectors (pTRBOcp) that enabled systemic movement of the virus-encoded constructs. pTRBOcp-*At*ORFs were agroinfiltrated (OD_600_ = 0.5) into the third youngest leaf (deemed L0) of individual 4-week-old *N. benthamiana* plants. The ORF-containing virus spread throughout the plant for 13 days, at which point the fourth leaf above L0 (deemed L4) was harvested for endogenous transcript (qPCR) analysis.

### Quantitative PCR (qPCR) for Synthetic Reporter and Endogenous Transcript Analysis

#### Primer design for detecting endogenous N. benthamiana transcripts

Sol Genomics Network nucleotide BLAST tool was used to query coding sequences of *Arabidopsis thaliana* type-A response regulators (RRs) for *N. benthamiana* homologs. Within the region of greatest homology in the top *N. benthamiana* candidate, primers (∼20nt in length; annealing temp: 59-60 °C) were designed to amplify an ∼120-150nt region. The NCBI primer design tool (https://www.ncbi.nlm.nih.gov/tools/primer-blast/) was used to avoid off-target binding and primer pair self-complementarity. See Supplementary Table 4 for qPCR primers used.

#### qPCR

First-strand cDNA (∼50 ng/μL) was diluted 1:5 to ensure the sample concentration was in the linear range of the primer set efficiency^61^. Diluted cDNA mixed with the appropriate primers (final concentration: 0.25 mM) and SensiMix™ SYBR® Hi-ROX was PCR-amplified using an Applied Biosystems™ QuantStudio™ 6 Pro (ThermoFisher). *N. benthamiana* Protein Phosphatase 2A (PP2A) was used as a housekeeping gene^62^, and the 2^−ΔΔCT^ method^63^ was used to calculate transcript fold change values.

### Statistical Tests

Statistical tests were performed as described in figure captions using GraphPad Prism Version 10.4.2 (534). For unpaired, two-way *t*-tests, F test was used to ensure equal variances, except for Figure 5C, where discrepancy is reported instead. For ANOVA, Brown-Forsythe test was used to ensure there was no statistically significant differences in standard deviations amongst the tested groups. If variances were deemed equal, ordinary one-way ANOVA with Tukey’s multiple comparisons test was used. If variances were unequal, Welch’s ANOVA with Dunnett’s T3 multiple comparisons test was used. If data did not pass Pearson’s test for normality, Kruskal-Wallis ANOVA with Dunn’s multiple comparisons test was used. Additional parameters for all null hypothesis testing can be found in Supplementary Table 10.

---

**Extended Data Figure 1.**
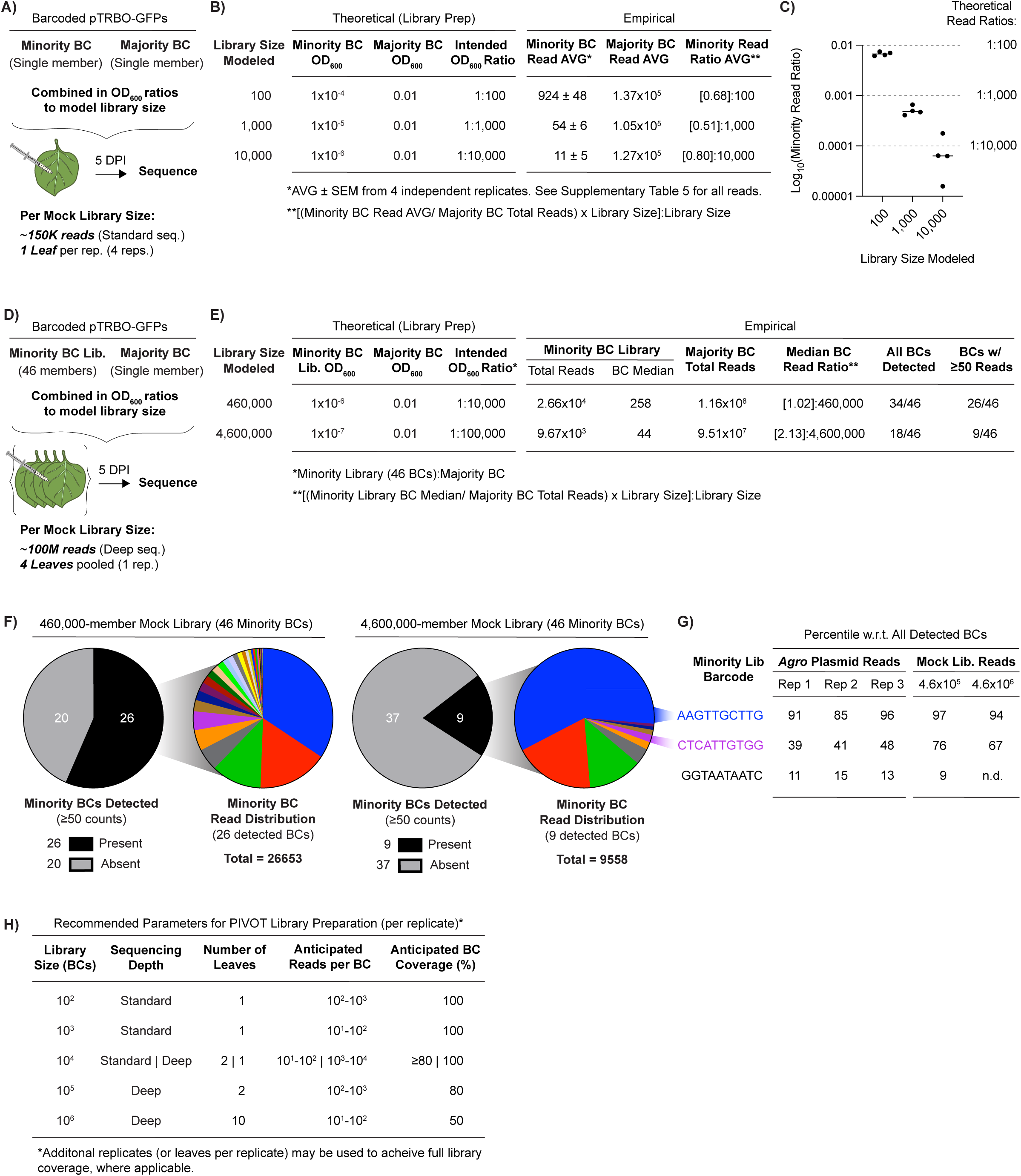
Mock libraries simulate upper limits for number of barcodes detectable in a single *N. benthamiana* leaf. (A) Workflow for mock libraries sizing 10^2^-10^4^ members: two *Agro* strains each harboring an independent barcoded pTRBO-GFP were combined in OD_600_ ratios described in (B) and infiltrated into one *N. benthamiana* leaf. At 5 DPI, four replicate samples were sent for standard depth sequencing (∼150,000 reads per sample). (B) Theoretical (left): OD_600_ for *Agro* strains harboring the minority and majority BCs to achieve the intended OD_600_ ratio modeling the indicated library size. Empirical (right): average read counts for minority and majority barcodes from four independent replicates (see Supplementary Table 5 for all read counts). (C) Median and four replicate log-transformed minority read ratios per mock library, calculated by dividing minority by majority barcode read count per replicate. Dotted lines indicate theoretical read ratios for respective library sizes (see Supplementary Note 1). (D) Workflow for mock libraries sizing 10^5^-10^6^ members: two *Agro* strains – one harboring a 46-member barcoded pTRBO-GFP library (Minority BC Lib.) and another harboring an independent barcoded pTRBO-GFP (Majority BC) – were combined in OD_600_ ratios described in (E) and infiltrated into four *N. benthamiana* leaves. At 5 DPI, tissue from four leaves was combined into a single sample and sent for deep sequencing (∼100,000,000 reads per sample). (E) Theoretical (left): OD_600_ for *Agro* strains harboring the minority library and the majority BC to achieve the intended OD_600_ ratio modeling the indicated library size. Empirical (right): counts for the minority library (total reads and median for a single BC) and majority barcodes (total reads) from one replicate; tabulated barcodes detected (total and exceeding a 50 read count threshold). (F) Pie charts detailing the number of barcodes within the 46-member minority BC library with ≥50 reads in the 460,000 and 4,600,000-member mock libraries (greyscale; left) and the read distribution for individual BCs within the detected cohort (colored; right). (G) Tabulated barcode representation percentiles for select members of the minority BC library across three independent replicate *Agro* plasmid and two *in planta* sequencing events. Barcode color corresponds to slice color in right pie chart of the 4,600,000-member mock library in (F); bottom barcode (black) not detected (n.d.) in the largest library tested. Percentiles calculated as [number of values ranked below BC/ total BCs detected] multiplied by 100. See Supplementary Table 5 for source reads. (H) Summarized recommendations (per replicate) for PIVOT library preparation in *N. benthamiana*, based on empirical data from (B) and (E).

**Extended Data Figure 2.**
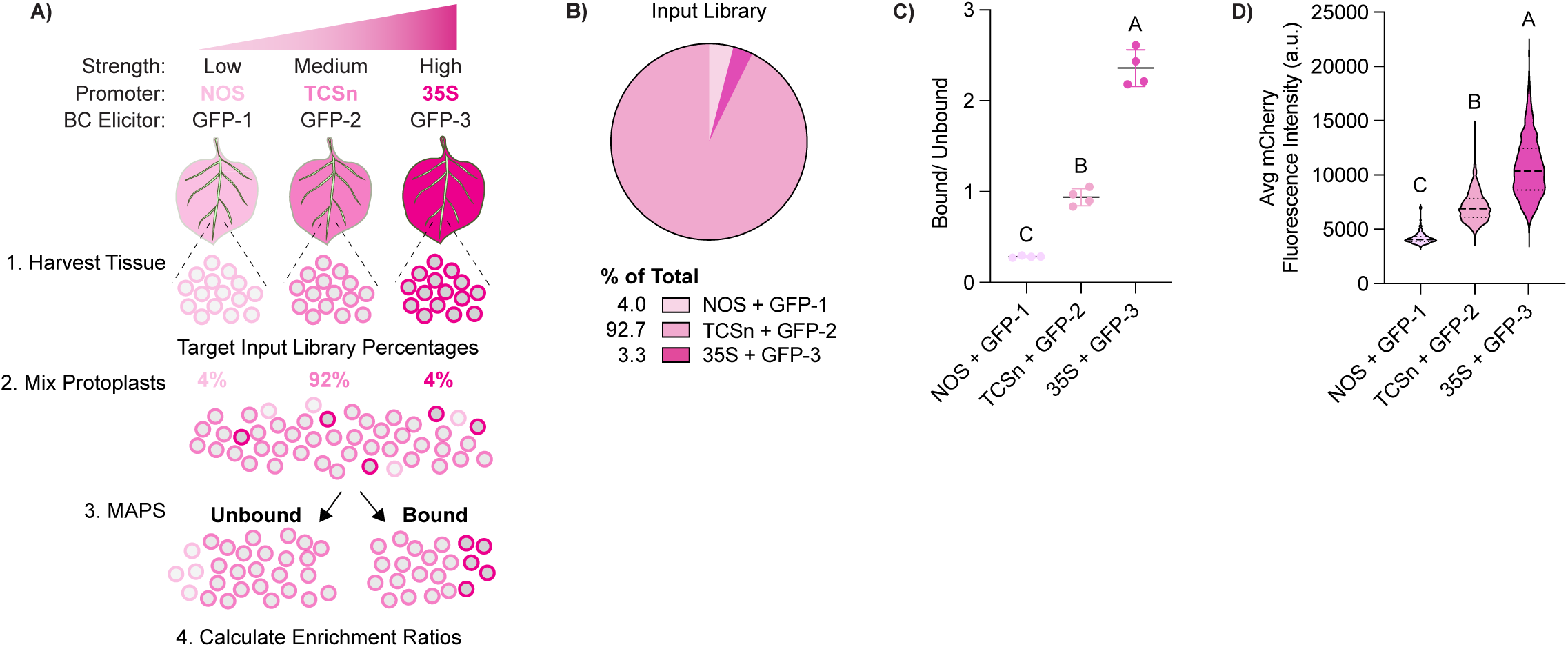
MAPS distinguishes protoplasts by reporter expression level. (A) Workflow for assessing MAPS ability to separate a protoplast population with varied reporter expression: independent leaves were co-infiltrated with HaloTag-mCherry driven by promoters of increasing strength (low: NOS (nopaline synthase promoter); medium: TCSn; and high: 35S) together with an identifying barcoded pTRBO-GFP. Protoplasts from each promoter cohort were mixed and then separated using MAPS. (B) Composition of the mixed protoplast library before MAPS separation (Input), quantified by barcode sequencing (average of four replicates). (C) Mean and standard deviation MAPS enrichment ratios (Bound/Unbound) for each promoter-GFP pairing from four independent replicates. See Supplementary Table 7 for read counts used to calculate enrichment ratios. Letters indicate statistically significant groups determined by a Brown-Forsythe and Welch ANOVA. (D) Violin plots representing the average mCherry fluorescence per cell for promoter cohorts, calculated from Cytation 5 fluorescence images. Median and quartiles indicated by dotted lines. Letters indicate statistically significant groups determined by a Kruskal-Wallis test. NOS: n = 164; TCSn: n = 972; 35S: n = 1215.

**Extended Data Figure 3.**
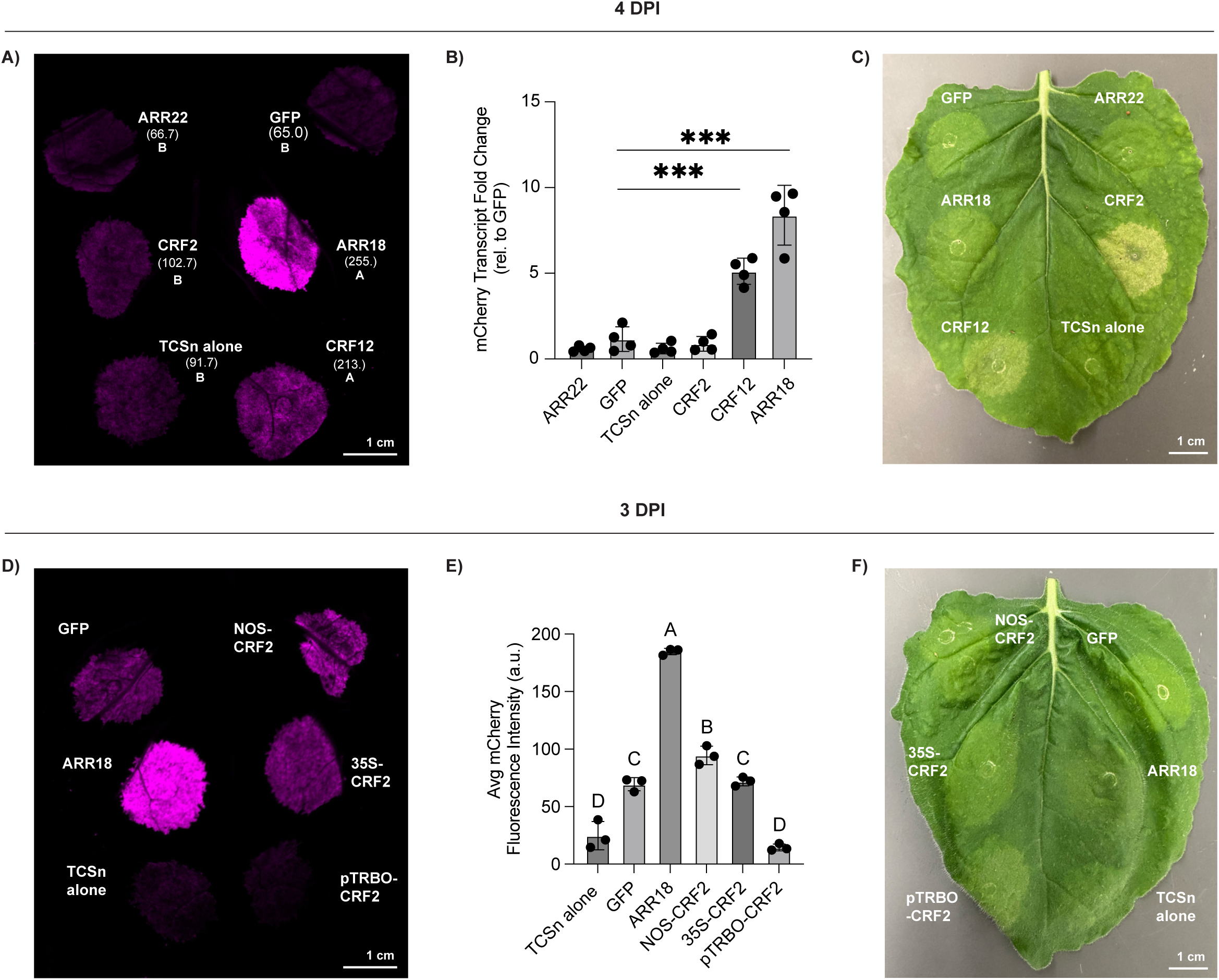
Post-screen assessment of CRF2 and CRF12-mediated TCSn activation. (A) iBright image of mCherry fluorescence four DPI for leaf spots co-infiltrated with a pTRBO-encoded screen candidate (OD_600_ = 0.1) and TCSn-mCherry reporter (OD_600_ = 0.5); abaxial leaf shown. Fluorescence values represent the mean of three repeat measures calculated using ImageJ. Letters signify statistically significant groups from Welch’s ANOVA with Dunnett’s T3 multiple comparisons test. (B) Quantification of TCSn-mCherry reporter transcript in tissue harvested from leaf spots shown in (A). Transcript quantifications normalized to a GFP control. Mean and standard deviation from 4 replicates plotted. Asterisks indicate significance values for unpaired two-tailed *t*-tests between individual query and GFP, *** p < 0.001. (C) Image of adaxial leaf in (A) indicating CRF-induced cell death. (D) iBright image of mCherry fluorescence three DPI for leaf spots co-infiltrated with a pEAQ-encoded screen candidate (except pTRBO-CRF2) at (OD_600_ = 0.1) and TCSn-mCherry reporter (OD_600_ = 0.5). Promoters of varied strength (NOS: low; 35S: medium; double 35S /pTRBO: high) control CRF2 expression to the right of the abaxial midrib. (E) Mean and standard deviation mCherry fluorescence calculated using ImageJ from three replicate leaves infiltrated as shown in (D). Letters indicate statistically significant groups from an ordinary one-way ANOVA with Tukey’s multiple comparisons test. (F) Image of adaxial leaf in (D) indicates less CRF-induced cell death at an earlier timepoint and using lower-strength promoters.

