## Supplementary Figures and Information for "An *in planta* single-cell screen to accelerate functional genetics"

**Table of Contents**

Supplementary Figures 2

Supplementary Notes 14

Supplementary Methods 18

Supplementary References 19

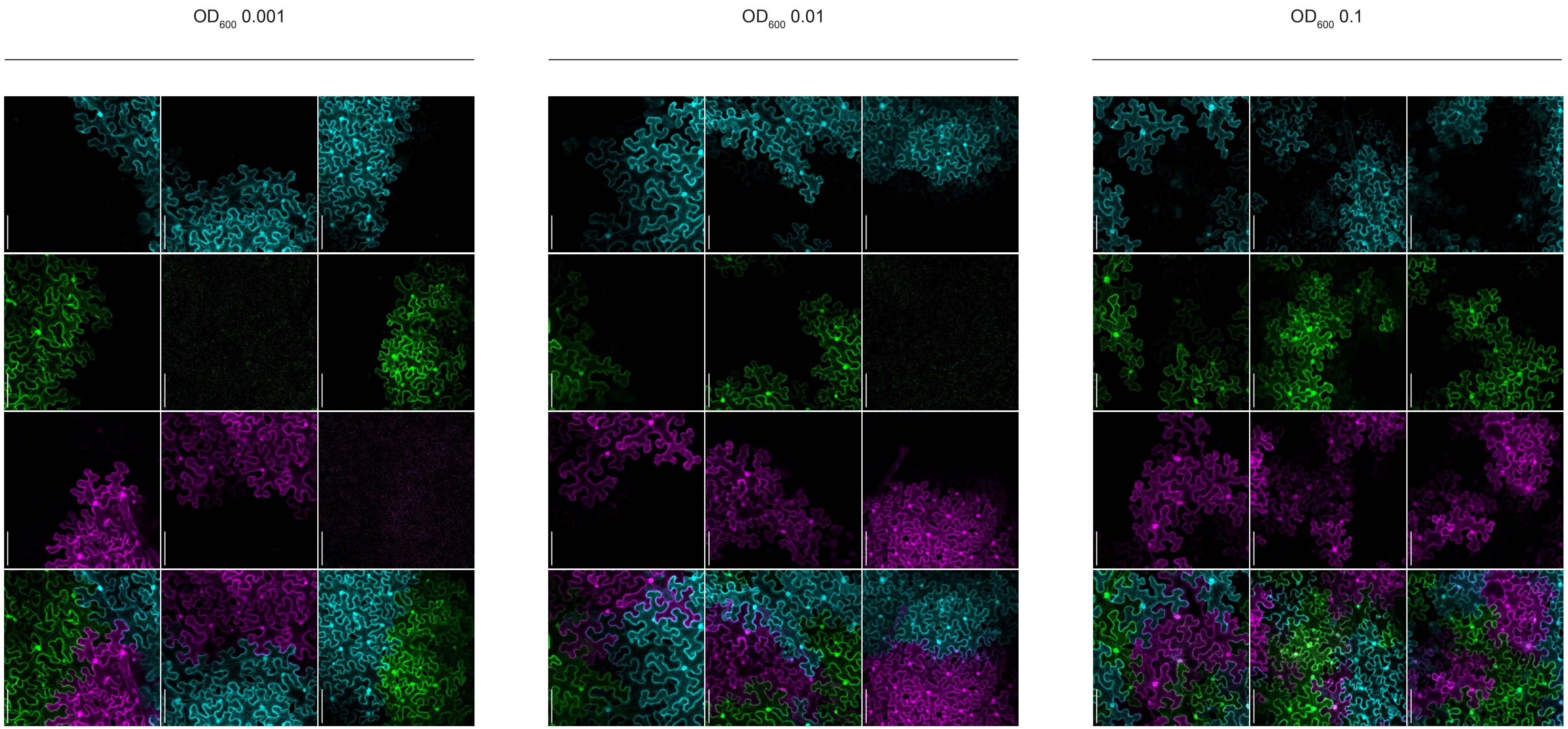
OD$_{600}$ 0.001
OD$_{600}$ 0.01
OD$_{600}$ 0.1

**Supplementary Figure 1. pTRBO vector exhibits superinfection exclusion and high tissue coverage regardless of *Agro* delivery $OD_{600}$.** Confocal microscopy images of *N. benthamiana* tissue five days post *Agro*infiltration with pooled pTRBO-encoded fluorescent proteins, as in main text Figure 2C. Three representative image series shown for each *Agro* delivery $OD_{600}$. Top, middle, and bottom rows within each $OD_{600}$ condition represent top (youngest), middle, and bottom leaves of 4-5-week-old *N. benthamiana* plants.

A)

| Construct | Intended Mock Library Ratio (mCherry:GFP) 1:100 | 1:1,000 | 1:10,000 | |
|---|---|---|---|---|
| pTRBO-mCherry | $1x10^{-4}$ | $1x10^{-5}$ | $1x10^{-6}$ | $OD_{600}$ |
| pTRBO-GFP | 0.01 | 0.01 | 0.01 | $OD_{600}$ |

B)

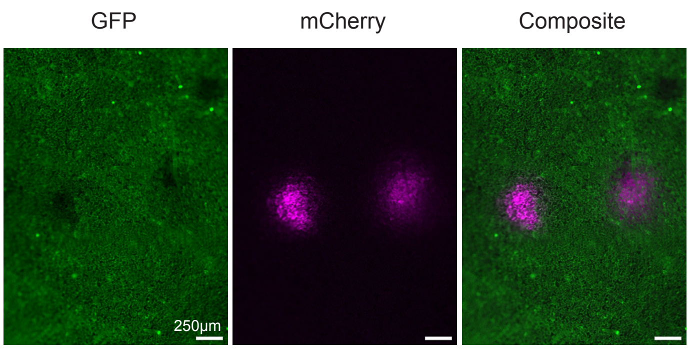


C)

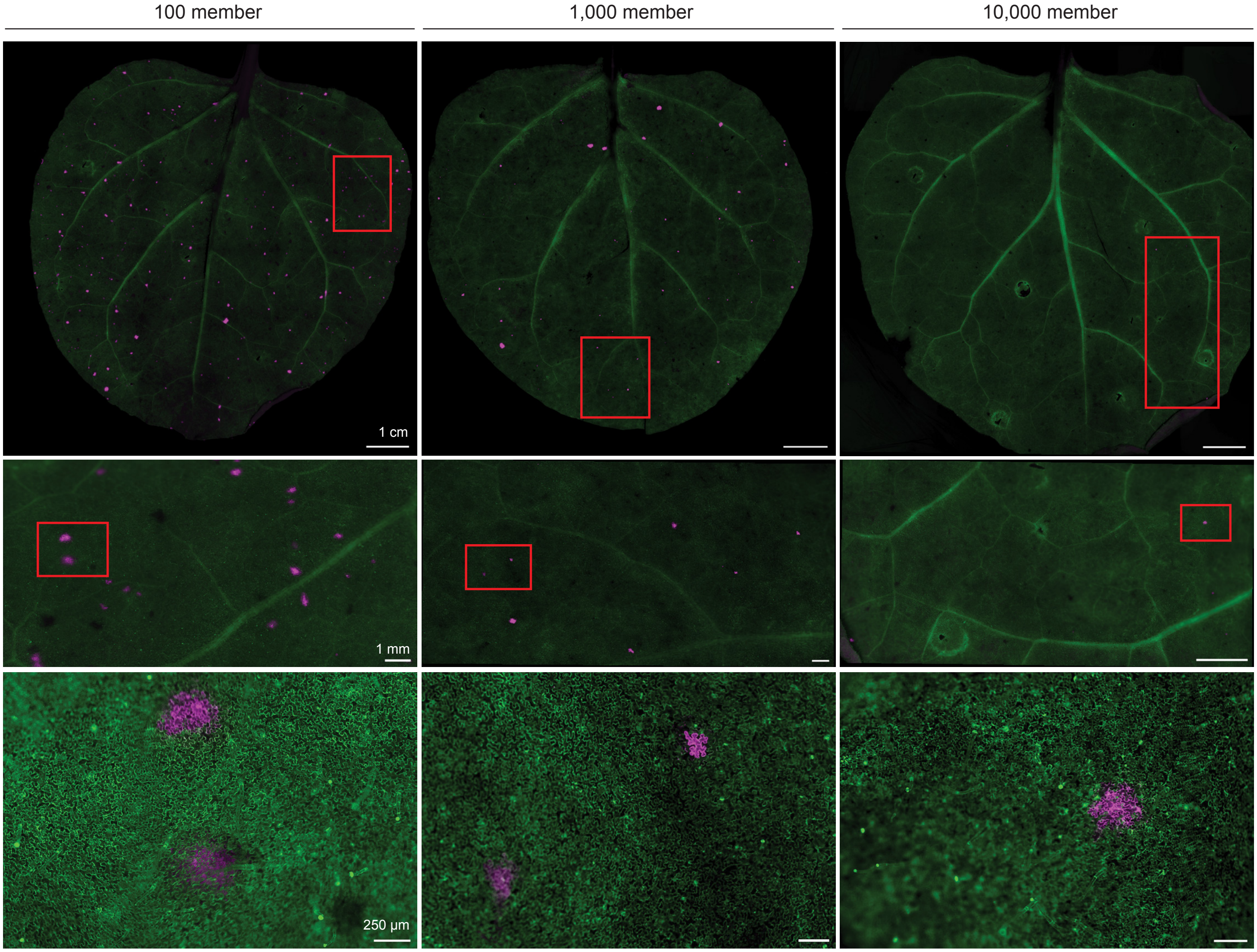


**Supplementary Figure 2. Viral delivery of mock gene libraries to *N. benthamiana*.** (A) *Agro* $OD_{600}$ values for pTRBO-mCherry (minority member) and pTRBO-GFP (bulk member) to achieve intented mock library ratios in (C) and main text Figure 2E. (B) Widefield stereoscope image of highest magnification frame in main text Figure 2E, with separate channels for GFP and mCherry shown to demonstrate superinfection exclusion at the cellular level during pooled pTRBO-FP delivery. (C) Widefield stereoscope images of *N. benthamiana* leaves five DPI with mock libraries. Red boxes indicate the magnification location for the next panel below within the same column.

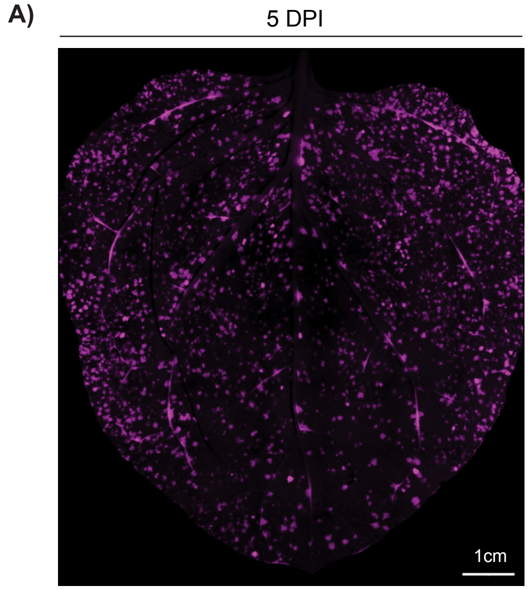


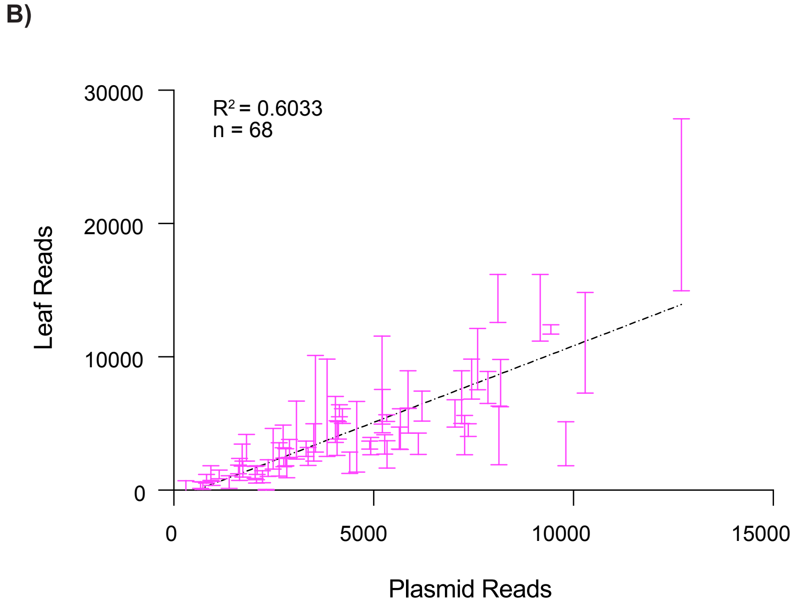


**Supplementary Figure 3. Quantification of in-leaf barcode representation for equal-length pTRBO-mCherry library.** (A) Widefield stereoscope image of *N. benthamiana* leaf infiltrated five DPI with 68-member pTRBO-mCherry-GUS library. The RNA insert size in the pTRBO vector totaled ~2.5 kb. (B) Plot of barcode reads before infiltration (*Agro* Plasmid) and five DPI (Leaf cDNA). Each mark represents the standard deviation of 4 replicate reads for an individual pTRBO-mCherry barcode, n = 68. Dotted line indicates linear regression, $R^2$ = 0.6033.

A)

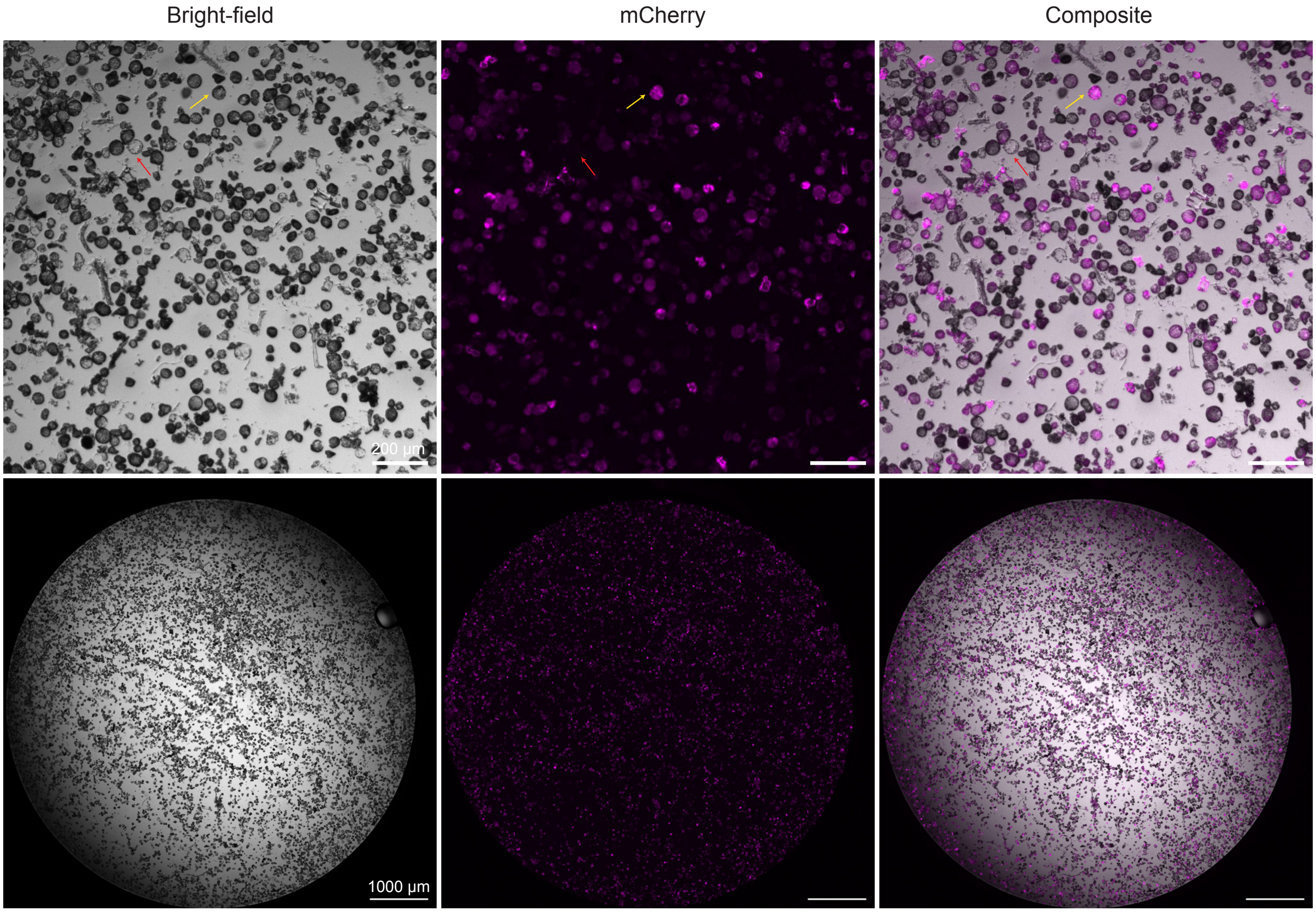


B)

**Cell Counts**

| | Rep 1 | Rep 2 | Rep 3 |
|---|---|---|---|
| Bright-field | 4207 | 4090 | 4304 |
| mCherry | 3986 | 3904 | 4059 |
| % Transformed | 94.7 | 95.5 | 94.3 |

C) **Estimate for number of transformed, harvestable protoplasts per leaf**

Based on cell counts in (A)

Dilution in (A): **1/20**

Well size: **200 µL**

Transformed cells/well (B): **4000**

Leaf count: **1/2**

Harvest volume: **3 mL**

$$\frac{\text{cells}}{\text{mL}} = \frac{1}{\text{dilution}} \times \frac{1}{\text{well size}} \times \frac{\text{cells}}{\text{well}} \times \frac{1000\ \text{uL}}{1\ \text{mL}}$$

$$\frac{\text{cells}}{\text{mL}} = 20 \times \frac{1}{200\ \text{uL}} \times 4000 \times \frac{1000\ \text{uL}}{1\ \text{mL}}$$

$$\frac{400{,}000\ \text{cells}}{\text{mL}} \times \frac{3\ \text{mL cells}}{1/2\ \text{leaf}} = \mathbf{2.4 \times 10^{6}\ \text{cells/ leaf}}$$

**Supplementary Figure 4. Image analysis of intact, transformed single cells following protoplasting.** (A) Images of protoplasts harvested from *N. benthamiana* tissue infiltrated with 35S-HaloTag-mCherry, zoomed (top) and whole-well (bottom). Yellow and red arrows indicate mCherry-positive and mCherry-null prototoplasts, respectively. (B) Cell counts derived from images in (A), bottom. All cells were gated on >0.6 circularity and >5000 a.u. for bright-field intensity. A second round of gating (>2000 a.u., mCherry) differentiated mCherry-positive from mCherry-null cells. Cell count gating was performed using Cytation 5 Gen5 software. (C) Estimated number of transformed, harvestable protoplasts per leaf based on preparation of sample imaged in (A) and cell counts in (B).

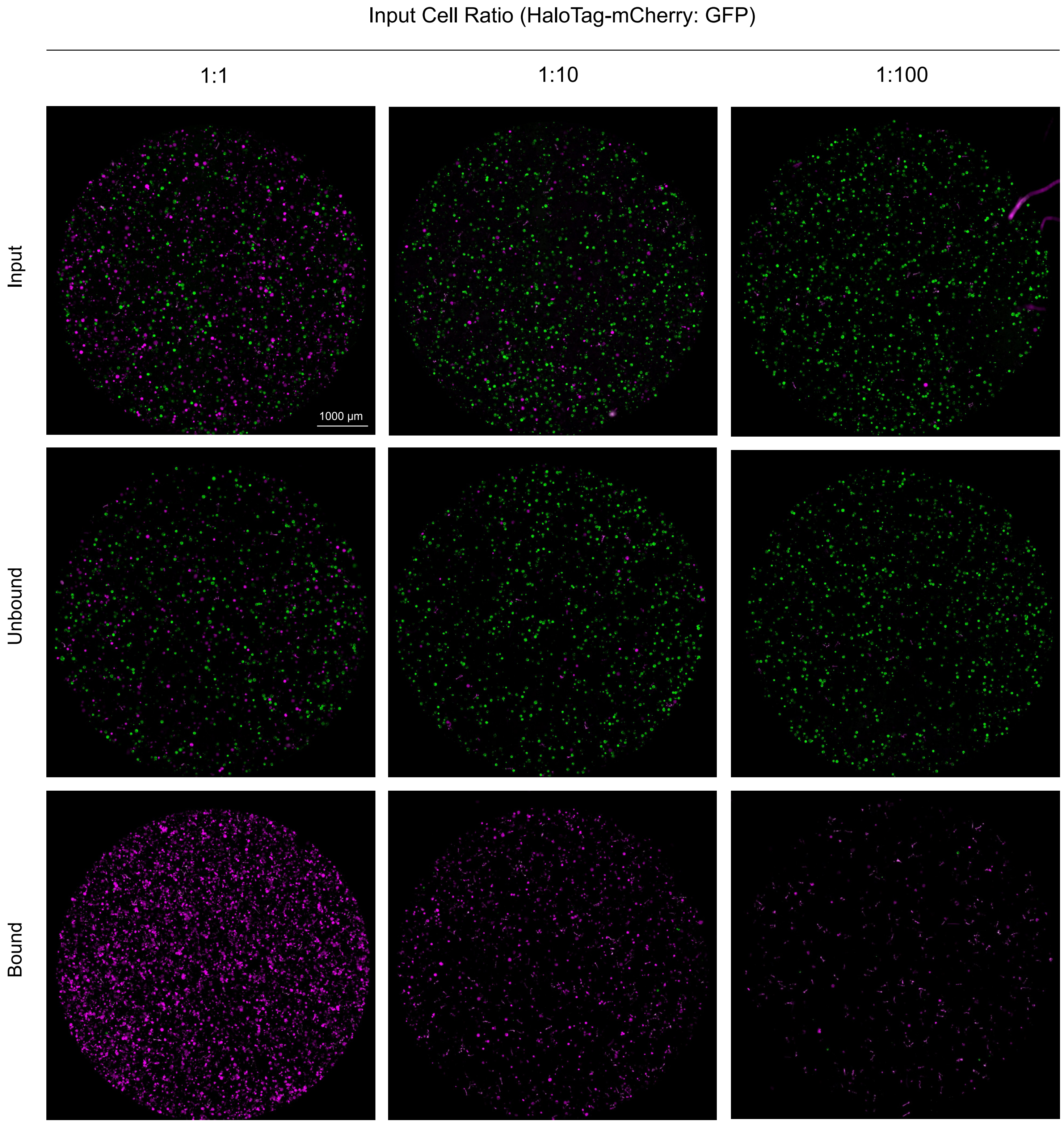


**Supplementary Figure 5. MAPS separation of mixed protoplast population.** Whole-well fluorescence microscopy images of mixed protoplast populations before (Input) and after (Unbound, Bound) MAPS. Source images for Figure 4D.

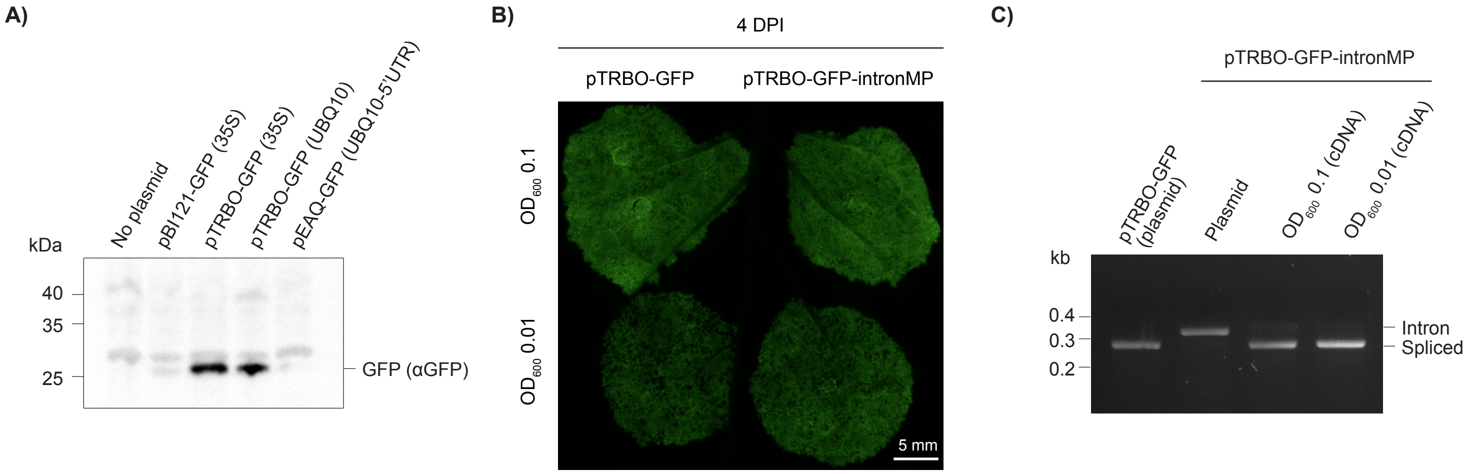


**Supplementary Figure 6. Vector intron does not disturb *in planta* function of pTRBO-delivered cargo.** (A) Western blot (anti-GFP) of *Agrobacterium* lysates transformed with various vector-promoter pairs driving GFP expression. (B) Leaf spot image demonstrating GFP fluorescence in *N. benthamiana* four DPI with regular pTRBO-GFP or pTRBO-GFP with an intron in the TMV movement protein (pTRBO-GFP-intronMP). (C) DNA gel of pre-infiltration plasmid or cDNA harvested from the respective leaf spot in (B).

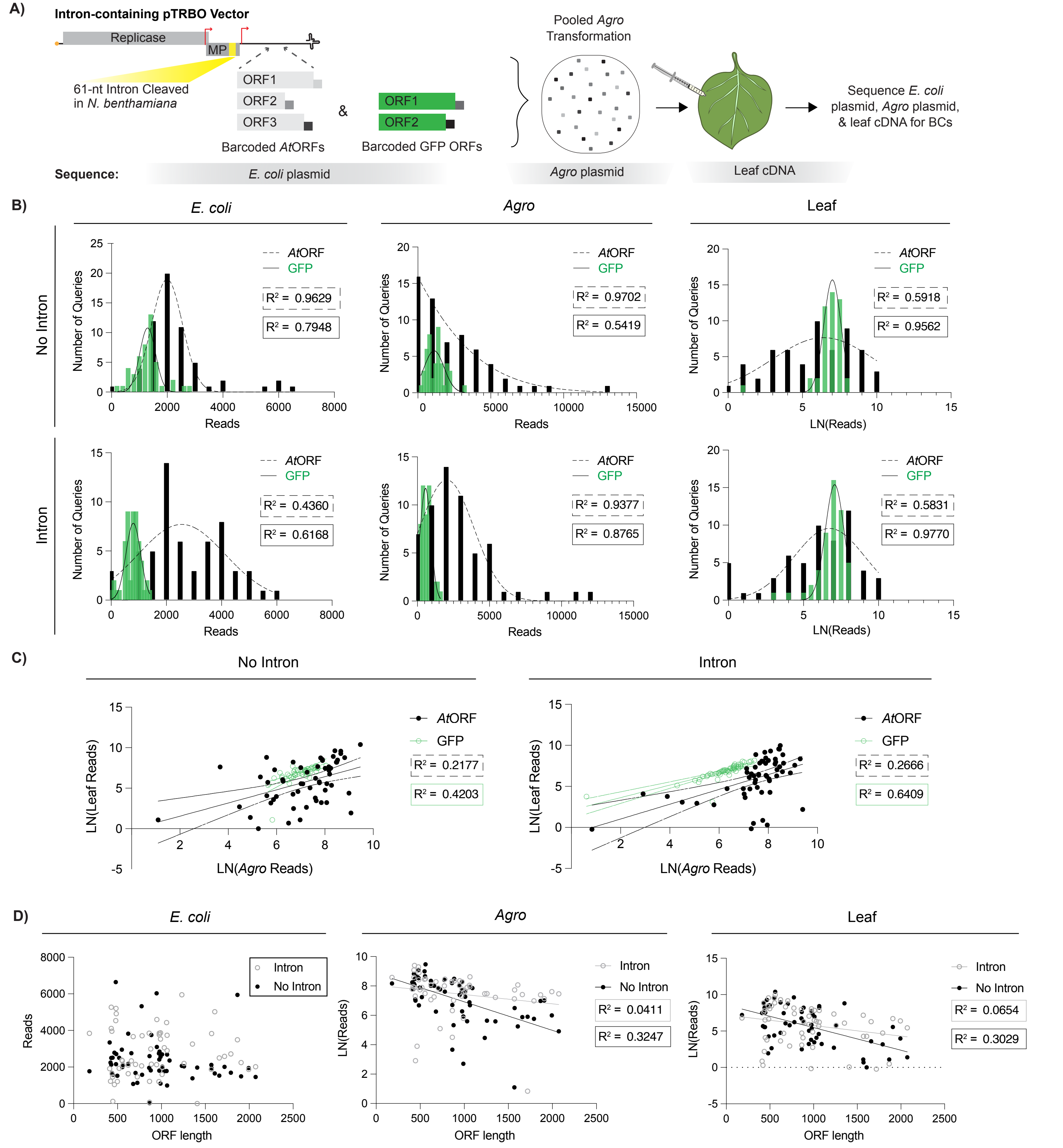


**Supplementary Figure 7. Intron-containing pTRBO vector corrects ORF toxicity in *Agrobacterium*.** (A) Workflow for assessing *At*ORF library representation dynamics: barcoded *At*ORFs and GFPs were cloned into intron-containing or non-intron-containing pTRBO vectors prior to pooled *Agro* transformation and leaf delivery. Pooled *E. coli* plasmid (one replicate), *Agro* plasmid (one replicate), and leaf cDNA (four replicates) were sequenced for individual barcodes. (B) Histograms comparing *At*ORF (black) and GFP (green) barcodes from *E. coli* plasmid, *Agro* plasmid, and leaf cDNA for ORF pools delivered with intron or no intron pTRBO vectors. Histograms fit with Gaussian curves. (C) Plots comparing *Agro* and leaf reads for pooled ORF libraries delivered from intron or non-intron-containing pTRBO vectors. Line of best fit with 95% confidence intervals plotted. (D) Linear plots from across library preparation stages comparing ORF size and read number for *At*ORFs delivered from intron or non-intron-containing pTRBO vectors. (B-D) See plots for linear fit $R^2$ values. For the non-intron library, n = 46 GFP and n = 58 for *At* ORFs; for the intron library, n = 54 GFP and n = 57 *At* ORFs.

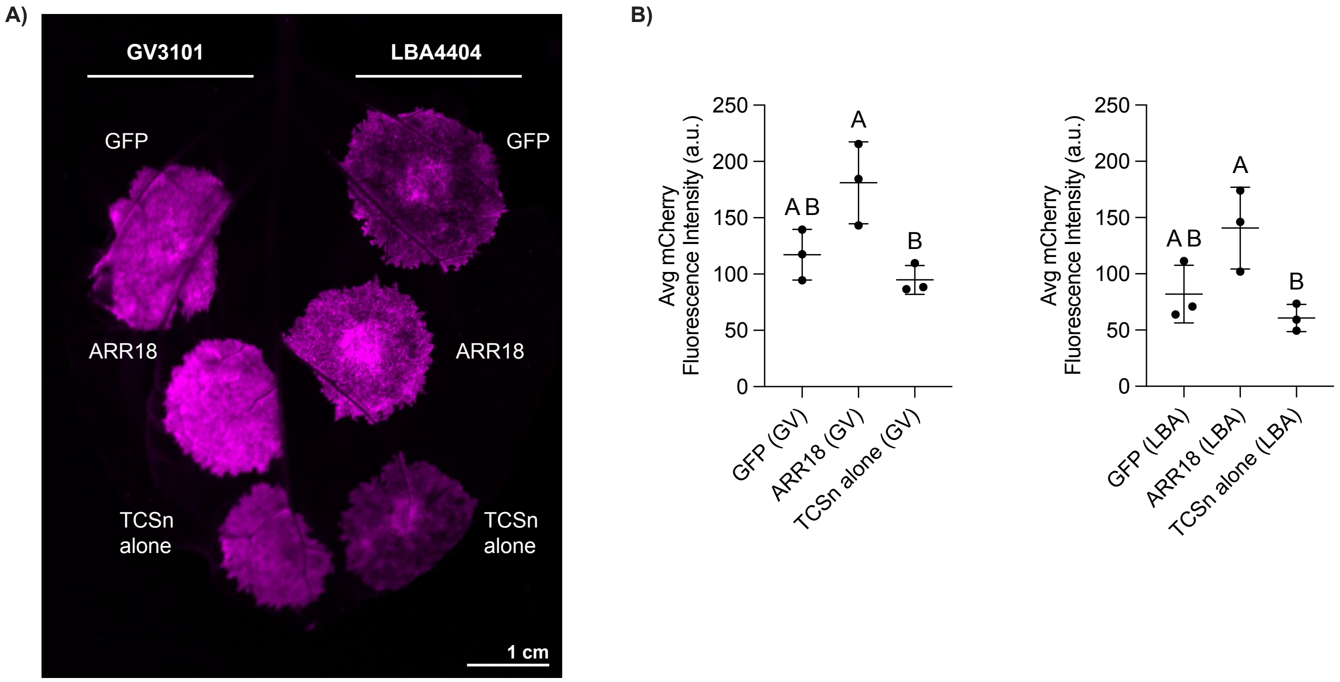


**Supplementary Figure 8. Relative ORF performance is not affected by the *Agrobacterium tumefaciens* strain used for delivery.** (A) iBright image of mCherry fluorescence three DPI for leaf spots co-infiltrated with a pTRBO-encoded screen candidate ($OD_{600}$ = 0.1) and TCSn-mCherry reporter ($OD_{600}$ = 0.5). Candidate and reporter constructs delivered to the leaf using the *tzs*-containing (cytokinin-producing) *Agro* strain GV3101 to the left of the abaxial midrib and the non-*tzs*-containing strain LBA4404 to the right. (B) Mean and standard deviation mCherry fluorescence calculated using ImageJ from three replicate leaves infiltrated as shown in (A). Letters indicate statistically significant groups from an ordinary one-way ANOVA with Tukey's multiple comparisons test. Right and left graphs quantify mCherry fluorescence from GV3101- and LBA4404-delivered queries in (A), respectively.

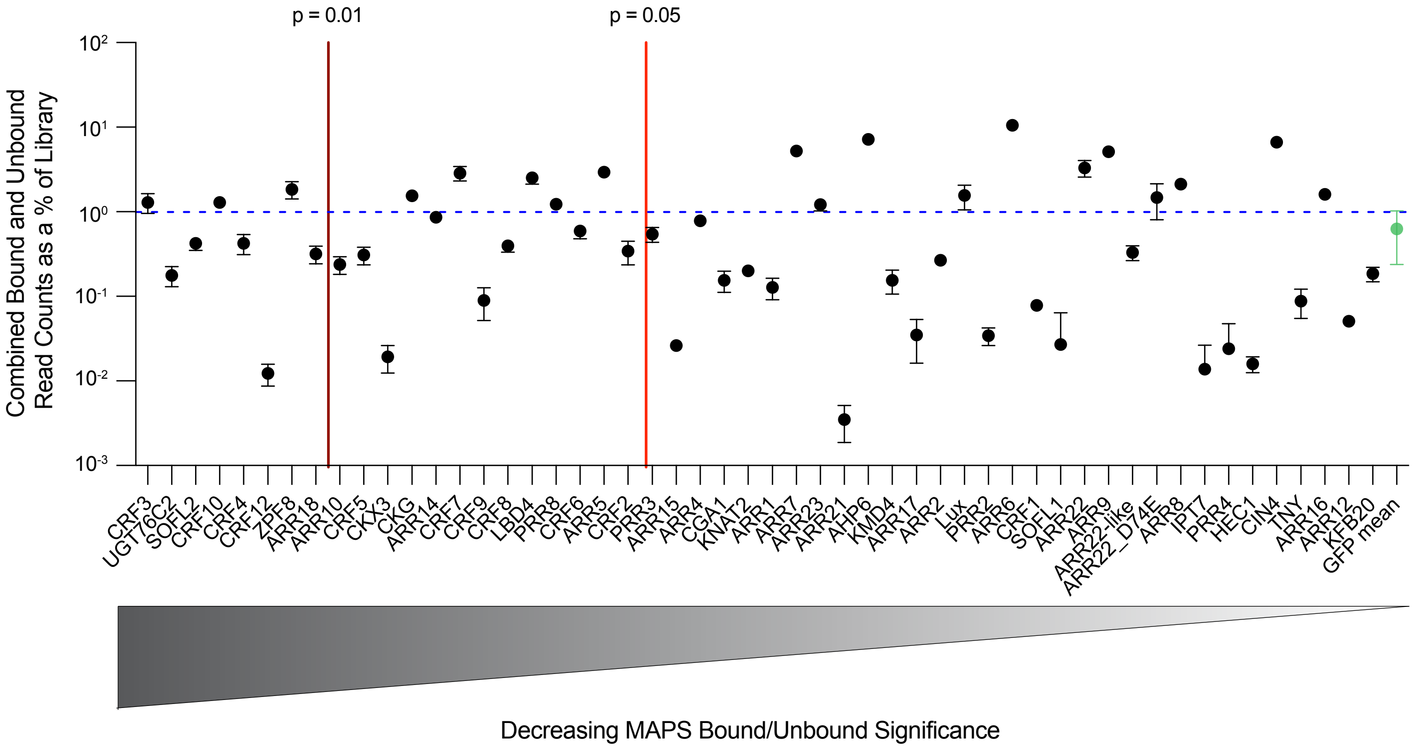


**Supplementary Figure 9. Representation of individual screen candidates in post-MAPS read counts.** Sum of Bound and Unbound read counts for each candidate as a percentatge of total MAPS Bound and Unbound read counts (each normalized to 150K reads, respectively) from the PIVOT screen (See Supplementary Table 9 for raw sequencing reads). The light and dark red lines denote p = 0.05 and p = 0.01, respectively, for significance of the MAPS Bound/Unbound enrichment score for an individual candidate compared to the GFP mean (See Supplementary Table 3 for p-values). Blue dotted line indicates the expected read count for a single query if all candidates were represented equally. Mean and standard deviation of four replicates plotted.

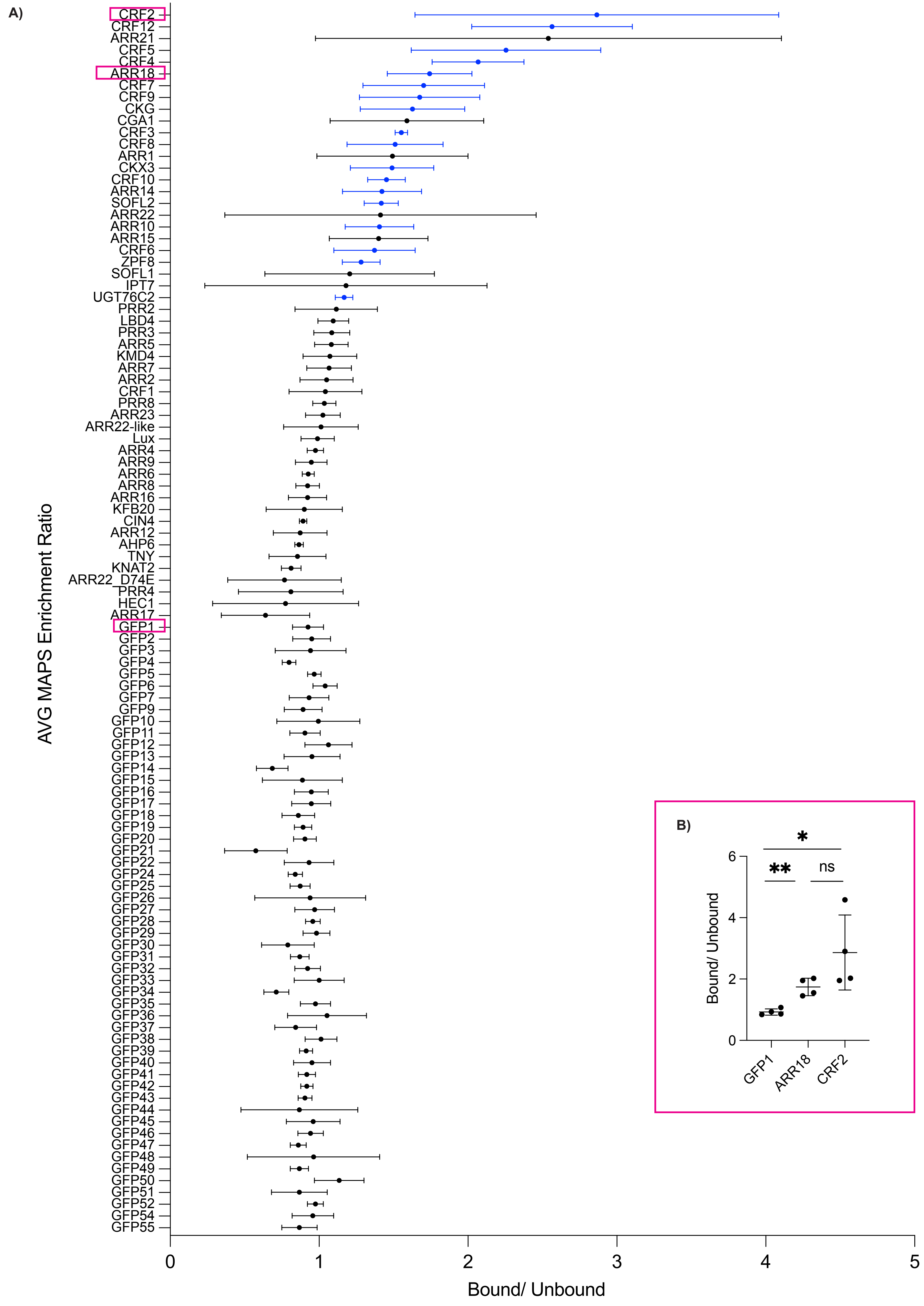

A)
AVG MAPS Enrichment Ratio
CRF2
CRF12
ARR21
CRF5
CRF4
ARR18
CRF7
CRF9
CKG
CGA1
CRF3
CRF8
ARR1
CKX3
CRF10
ARR14
SOFL2
ARR22
ARR10
ARR15
CRF6
ZPF8
SOFL1
IPT7
UGT76C2
PRR2
LBD4
PRR3
ARR5
KMD4
ARR7
ARR2
CRF1
PRR8
ARR23
ARR22-like
Lux
ARR4
ARR9
ARR6
ARR8
ARR16
KFB20
CIN4
ARR12
AHP6
TNY
KNAT2
ARR22_D74E
PRR4
HEC1
ARR17
GFP1
GFP2
GFP3
GFP4
GFP5
GFP6
GFP7
GFP9
GFP10
GFP11
GFP12
GFP13
GFP14
GFP15
GFP16
GFP17
GFP18
GFP19
GFP20
GFP21
GFP22
GFP24
GFP25
GFP26
GFP27
GFP28
GFP29
GFP30
GFP31
GFP32
GFP33
GFP34
GFP35
GFP36
GFP37
GFP38
GFP39
GFP40
GFP41
GFP42
GFP43
GFP44
GFP45
GFP46
GFP47
GFP48
GFP49
GFP50
GFP51
GFP52
GFP54
GFP55
0
1
2
3
4
5
Bound/ Unbound
B)
*
**
ns
6
4
2
0
Bound/ Unbound
GFP1
ARR18
CRF2

**Supplementary Figure 10. MAPS enrichment ratios for all PIVOT screen candidates.** (A) Mean and standard deviation of four replicate MAPS bound/unbound enrichment ratios for all PIVOT screen candidates. *At*ORF candidates ranked by decreasing average MAPS enrichment ratio from top to bottom; GFP population ordered numerically. Blue standard deviation lines indicate screen candidates yielding average enrichment ratios above 1 standard deviation of the GFP mean enrichment ratio and with $p < 0.05$. Pink outline indicates candidates selected for additional statistical analysis in (B) insert. (B) Mean and standard deviation of four replicate MAPS bound/unbound enrichment ratios for three screen candidates: two screen hits (CRF2, ARR18) and the median GFP candidate (GFP1). Asterisks indicate significance for two-tailed, unpaired Welch's t-tests between individual query and GFP, * $p < 0.05$ and ** $p < 0.01$.

**Supplementary Note 1**

We used amplicon sequencing to quantify representation of two differentially barcoded pTRBO-GFPs that were delivered to a leaf in *Agro* $OD_{600}$ ratios to model a 100-member library (i.e., minority BC: final $OD_{600}$ $1x10^{-4}$; majority BC: final $OD_{600}$ 0.01; Extended Data Figure 1A, B). For this mock 100-member library, the minority read ratio (calculated as [the minority barcode read count divided by the majority barcode read count multiplied by the intended library size]:intended library size) ranged between [0.61-0.74]:100 across four replicates, which is ~30% less than the theoretical minority read ratio of [1]:100 for a library of this size. We repeated this experiment for mock 1,000 and 10,000-member libraries, and achieved similar results: for each mock library, the average minority read ratio was slightly below, but within range of the expected ratio. Experimental error during serial dilution (i.e. a smaller than intended portion of the *Agro* population successfully transferred in successive serial dilutions) could account for the observed ~20-50% drop in observed compared to theoretical read ratio (e.g. instead of 1 in 100 (theoretical), 0.68 in 100 (observed); Extended Data Figure 1B, C). In sum, we suspect the slightly reduced experimental read count is an artifact of how we prepared the mock libraries in this experiment, and we anticipate this error could be avoided in future PIVOT iterations which implement pooled cloning methods rather than *Agro* strain mixing during library preparation.

For the experiments in Extended Data Figure 1A-C, we used a sequencing service that typically yields between 100,000-200,000 reads. Thus, we expected to near the limit of detection at this sequencing depth for the mock 10,000-member library, and indeed, we detected an average ± standard error of the mean (SEM) of 11 ± 5 minority barcode reads per replicate (Extended Data Figure 1B, Supplementary Table 5). Therefore, to truly test the limit of how many library members (barcodes) we could deliver and detect in a single (or few) leaves, we used a deeper sequencing method (~$10^2$ million reads per sample). For the 1:10,000 and 1:100,000 $OD_{600}$ ratio mock library experiments, the minority population was comprised of 46 barcoded GFP clones. Therefore, a single barcode represents 1 member of mock 460,000 and 4,600,000-member libraries, respectively. We infiltrated 4 leaves per ratio and pooled RNA from all 4 leaves to send for sequencing (Extended Data Figure 1D, E).

For the mock 460,000-member library, 32 out of 46 barcoded minority library members were detected, with 26 barcodes (57%) yielding over a 50-read count threshold; the number of total detected minority library barcodes decreased by about half to 17 members in the mock 4.6-million-member library, with 9 barcodes (20%) surpassing the 50-read count threshold (Extended Data Figure 1E, F). The number of reads per individual barcode within the minority library ranged from single digits to $10^3$ reads (out of a total ~$10^8$ reads for minority library and majority reads combined): a majority of detected members yielded on the order of $10^2$ reads in the 460,000-member library and $10^1$ reads in the 4.6-million-member library, respectively (Extended Data Figure 1E, Supplementary Table 5). Our previous data (Figure 3C, F and Supplementary Figure 3B) demonstrate that this spread in leaf reads for individual barcodes is expected given distribution of *Agro* plasmid reads within the 46-member minority barcoded library (Supplementary Table 5). Indeed, the most highly represented barcode in the mock

460,000 and 4,600,000-member libraries (AAGTTGCTTG) consistently ranked between the 85-96th percentile in *Agro* plasmid reads, while a barcode (GGTAATAATC) ranking in the 11-15th percentile in *Agro* plasmid reads placed in the 9th percentile for the mock 460,000 library and was not detected in the 4,600,000-member library, respectively (Extended Data Figure 1G).

Taken together, these data suggest that standard depth (~$10^5$ reads) and deep (~$10^2$ million reads) sequencing can readily detect $10^3$ and $10^4$ barcodes, respectively, from a single *N. benthamiana* leaf. We recommend using multiple leaves and/or multiple replicates to ensure adequate library coverage for $10^{5+}$ barcodes (e.g.1-2 leaves per $1x10^5$ barcodes to achieve an estimated ≥ 80% library coverage; Extended Data Figure 1H). These estimates for the number of barcodes we can reliably detect per leaf ($10^4$-$10^5$) seem in congruence with the number of harvestable, transformed protoplasts per leaf, which we estimate to be on the order of $10^6$ cells (Supplementary Figure 4C).

**Supplementary Note 2**

When we initially cloned individual *Arabidopsis* ORFs into the pTRBO vector, we noticed several constructs were unusually difficult to grow in Agro. We suspected this toxicity was caused by leaky expression of T-DNA in *Agro* (Supplementary Figure 6A). Thus, we decided to implement an intron in the pTRBO vector backbone to decrease expression in *Agro*. We inserted a *Agro*-derived 61-nucleotide intron (previously characterized in our lab; see main text reference 29) into the C-terminus of the TMV movement protein. We found that the intron was spliced out in *N. benthamiana* (Supplementary Figure 6C) and did not disrupt *in planta* activity of the pTRBO-delivered cargo (Supplementary Figure 6B).

In preparation for our pooled ORF screen, we wanted to assess the representation of biologically functional ORFs across stages of library preparation. We cloned our barcoded *At*ORFs and GFP ORFs into both a regular (no intron) and an intron-containing pTRBO-vector and combined equal plasmid ratios of the 112-member barcoded library for pooled transformation into *Agro* and delivery into *N. benthamiana*. We used amplicon sequencing to track the number of ORF-specific barcodes from (1) the manually prepped plasmid pool ("*E. coli*), (2) plasmid prepped following pooled *Agro* transformation ("*Agro*"), and (3) cDNA harvested from infiltrated tissue ("Leaf") (Supplementary Figure 6A).

Histograms plotting the *E. coli* read counts for the AtORF and GFP libraries in both the no-intron and intron-containing pTRBO vectors adopt normal Gaussian distributions (Supplementary Figure 7B, *E. coli*). Similarly, *Agro* read counts for the GFP library in no-intron and intron pTRBO vectors are normally distributed. Strikingly, *Agro* reads from the *At*ORF library in the no-intron vector are severely right-skewed; however, expressing this library in an intron-containing vector corrects the *Agro* read distribution to a normal Gaussian distribution (Supplementary Figure 7B, *Agro*).

Although adding the intron into the pTRBO vector relieved ORF toxicity and corrected library distribution in *Agro*, the intron did not yield a significant difference in leaf distribution for our *At*ORF library. Regardless of intron presence, GFP leaf reads demonstrated tight Gaussian distributions, while the *At*ORF library displayed a wide and less significant Gaussian distribution (Supplementary Figure 7B, Leaf). These data suggest that regardless of vector intron presence, ORF biological function – e.g., protein toxicity – may further contribute to library skewing *in planta* and underrepresentation of some candidate genes. This hypothesis is supported by the data in Supplementary Figure 7 (panel C, Intron), wherein some candidates that were present in the *Agro* library dropped out after leaf infiltration, perhaps indicating that the encoded proteins resulted in cell death

In addition to ORF identity, we were interested in analyzing the impact of ORF size on candidate representation in our intron and no-intron pTRBO-*At*ORF libraries. In the manually prepared *E. coli* pools, ORF size did not have a significant effect on candidate representation in the intron or no-intron libraries (Supplementary Figure 7D, *E. coli*). However, the no-intron library demonstrated a negative correlation between *Agro* reads and ORF size – likely due to the aforementioned leaky ORF expression in *Agro* – which was negated (indicated by lower $R^2$ value) by implementing the pTRBO intron (Supplementary Figure 7D, *Agro*). Correcting candidate representation in *Agro* extended to leaf reads, where the negative correlation between ORF size and leaf reads in the no-intron cohort was relieved in the intron cohort. Further, larger ORF sizes had more in-leaf reads in the intron-containing library compared to the no-intron cohort (Supplementary Figure 7D, Leaf). Taken together, the data from Supplementary Figure 7 demonstrate that the intron-containing pTRBO construct can improve library representation in *Agro* by relieving toxic leaky expression.

**Supplementary Note 3**

Interestingly, while CRF2 had the largest MAPS enrichment ratio (Figure 5C) and was the strongest inducer of *N. benthamiana* type-A RR genes out of all screen candidate tested (Figure 5H), it initially failed to induce significant TCSn transcript levels in follow-up spot assays (Extended Data Figure 3A, B). However, in these experiments, we noticed that CRF2 overexpression in many adjacent cells caused extensive cell death, potentially confounding reporter signal (Extended Data Figure 3C). We resolved this incongruency between PIVOT screen and leaf spot assay results by reducing the strength of the promoter driving CRF2 expression, which lessened, but did not fully relieve CRF2-induced cell death (Extended Data Figure 3D-F). This promoter swap solution highlights that the pTRBO vector – which harbors a double 35S promoter – may require tuning depending on the predicted activity of ORF candidates or the context of a PIVOT screen. Taken together, our investigation of CRF2 highlights the power of the PIVOT screening approach, which may be more amenable to interrogate candidates that are cytotoxic (and therefore overlooked) when assessed individually under conditions of high local concentration.

**Supplementary Note 4**

For our proof-of-concept PIVOT screen, we used the following statistical boundaries to delineate screen "hits": (A) a bound/unbound ratio beyond 1 standard deviation of the GFP mean and (B) significant p-value ($p < 0.05$ or $p < 0.01$) compared to the control GFP population. A total of 18 ORFs fulfilled condition (A) – ten with a p-value between

0.05-0.01 and an additional eight ORFs yielded p-value significance beyond 0.01 (Figure 5C, Supplementary Table 10).

When we pursued a handful of these candidates in follow-up leaf spot assays, all but CRF3 exhibited activation above reporter alone and GFP controls (CRFs 4,5,10, and ARR18 – Figure 5D-F; CRFs 2,12 – Extended Data Figure 3). It is important to note that although PIVOT-predicted "hits" demonstrated significant reporter activation in the leaf spot assay, the PIVOT enrichment ratio magnitudes were not necessarily correlative of relative performance in secondary assays. Closer consideration of the standard deviations (SD) for each ORF enrichment ratio demonstrate that many of the top PIVOT hits may be statistically indistinguishable (overlapping SD error bars) (Supplementary Figure 10A). For example, the average and SD PIVOT enrichment ratios for CRF2 and ARR18 were 2.87 (+/-1.22) and 1.74 (+/-0.28), which are both statistically significant compared to the middle 50% GFP (GFP1: 0.93 +/- 0.11), but not statistically different from each other (Supplementary Figure 10B).

Taken together, the data from our proof-of-concept cytokinin screen was capable of generating a list of "hit" candidates that were statistically significant from the control GFP population and that could be prioritized for further testing; however, for this screen in particular, we do not have evidence that the PIVOT enrichment ratios alone could be used to statistically differentiate between "hit" candidates or definitively assign strength of their biological activity. We anticipate future iterations of PIVOT that include more gene candidates and/or a larger dynamic range in enrichment ratios could harbor enough statistical power to conclusively differentiate between "hit" candidates based on enrichment ratio. However, as is the case with many biological screens, here, further examination of individual candidates in secondary assays would be necessary to make claims of comparative function between candidates following preliminary screening.

### Supplementary Methods (Relevant to Supplementary Figure 6)

#### Western Blot

*Agrobacterium* cultures were pelleted by centrifugation at 6,000 × g for 5 minutes, washed twice with induction buffer (10 mM MES, pH 5.6, 10 mM $MgCl_2$), and resuspended. Optical density at 600 nm ($OD_{600}$) was measured, and cultures were normalized accordingly. The cells were then re-pelleted, resuspended in 2× LDS sample buffer containing 10% β-mercaptoethanol, and denatured at 75°C for 30 minutes. Soluble protein fractions were obtained by sonication and centrifugation and subjected to NuPAGE gel electrophoresis (ThermoFisher) at 200 V for 45 minutes using 1× MOPS running buffer.

Proteins were transferred to a PVDF membrane using the Trans-Blot Turbo system (Bio-Rad), following the manufacturer's protocol. The membrane was blocked in EveryBlot blocking buffer (Bio-Rad) for 20 minutes at room temperature, then incubated overnight at 4°C with chicken anti-GFP primary antibody (1:5000; Fisher Scientific, NC1666473) with gentle agitation. The next day, the membrane was washed three times for 10 minutes each with 1× TBST, then incubated for 1 hour at room temperature with goat anti-chicken secondary antibody (1:5000; Abcam, ab97135) in 1× TBST. After five 5-minute washes in 1× TBST and a final wash in 1× PBS, the membrane was developed using Clarity ECL substrate (Bio-Rad) for 5 minutes and imaged with an iBright FL1500 imager (ThermoFisher).

#### DNA Gel Electrophoresis

Plasmid purified from *E. coli* or cDNA prepared from harvested tissue after infiltration were loaded on a 0.8% agarose gel in TAE buffer. Samples were run on an Owl EasyCast Gel Electrophoresis System (ThermoFisher) at 150 V for 30 minutes.

(11) Horák, J.; Grefen, C.; Berendzen, K. W.; Hahn, A.; Stierhof, Y.-D.; Stadelhofer, B.; Stahl, M.; Koncz, C.; Harter, K. The Arabidopsis Thaliana Response Regulator

ARR22 Is a Putative AHP Phospho-Histidine Phosphatase Expressed in the Chalaza of Developing Seeds. *BMC Plant Biol.* **2008**, *8* (1), 77. https://doi.org/10.1186/1471-2229-8-77.
(12) Schaller, G. E.; Kieber, J. J.; Shiu, S.-H. Two-Component Signaling Elements and Histidyl-Aspartyl Phosphorelays†. *Arab. Book Am. Soc. Plant Biol.* **2008**, *6*, e0112. https://doi.org/10.1199/tab.0112.
(13) Gattolin, S.; Alandete-Saez, M.; Elliott, K.; Gonzalez-Carranza, Z.; Naomab, E.; Powell, C.; Roberts, J. A. Spatial and Temporal Expression of the Response Regulators ARR22 and ARR24 in Arabidopsis Thaliana. *J. Exp. Bot.* **2006**, *57* (15), 4225–4233. https://doi.org/10.1093/jxb/erl205.
(14) Park, J.; Lee, S.; Park, G.; Cho, H.; Choi, D.; Umeda, M.; Choi, Y.; Hwang, D.; Hwang, I. CYTOKININ-RESPONSIVE GROWTH REGULATOR Regulates Cell Expansion and Cytokinin-Mediated Cell Cycle Progression. *Plant Physiol.* **2021**, *186* (3), 1734–1746. https://doi.org/10.1093/plphys/kiab180.
(15) Bilyeu, K. D.; Cole, J. L.; Laskey, J. G.; Riekhof, W. R.; Esparza, T. J.; Kramer, M. D.; Morris, R. O. Molecular and Biochemical Characterization of a Cytokinin Oxidase from Maize1. *Plant Physiol.* **2001**, *125* (1), 378–386. https://doi.org/10.1104/pp.125.1.378.
(16) Rashotte, A. M.; Mason, M. G.; Hutchison, C. E.; Ferreira, F. J.; Schaller, G. E.; Kieber, J. J. A Subset of Arabidopsis AP2 Transcription Factors Mediates Cytokinin Responses in Concert with a Two-Component Pathway. *Proc. Natl. Acad. Sci.* **2006**, *103* (29), 11081–11085. https://doi.org/10.1073/pnas.0602038103.
(17) Cutcliffe, J. W.; Hellmann, E.; Heyl, A.; Rashotte, A. M. CRFs Form Protein-Protein Interactions with Each Other and with Members of the Cytokinin Signalling Pathway in Arabidopsis via the CRF Domain. *J. Exp. Bot.* **2011**, *62* (14), 4995–5002. https://doi.org/10.1093/jxb/err199.
(18) Swinka, C.; Hellmann, E.; Zwack, P.; Banda, R.; Rashotte, A. M.; Heyl, A. Cytokinin Response Factor 9 Represses Cytokinin Responses in Flower Development. *Int. J. Mol. Sci.* **2023**, *24* (5), 4380. https://doi.org/10.3390/ijms24054380.
(19) Zhang, J.; Vankova, R.; Malbeck, J.; Dobrev, P. I.; Xu, Y.; Chong, K.; Neff, M. M. AtSOFL1 and AtSOFL2 Act Redundantly as Positive Modulators of the Endogenous Content of Specific Cytokinins in Arabidopsis. *PLOS ONE* **2009**, *4* (12), e8236. https://doi.org/10.1371/journal.pone.0008236.
(20) Hass, C.; Lohrmann, J.; Albrecht, V.; Sweere, U.; Hummel, F.; Yoo, S. D.; Hwang, I.; Zhu, T.; Schäfer, E.; Kudla, J.; Harter, K. The Response Regulator 2 Mediates Ethylene Signalling and Hormone Signal Integration in Arabidopsis. *EMBO J.* **2004**, *23* (16), 3290–3302. https://doi.org/10.1038/sj.emboj.7600337.
(21) Chiang, Y.-H.; Zubo, Y. O.; Tapken, W.; Kim, H. J.; Lavanway, A. M.; Howard, L.; Pilon, M.; Kieber, J. J.; Schaller, G. E. Functional Characterization of the GATA Transcription Factors GNC and CGA1 Reveals Their Key Role in Chloroplast Development, Growth, and Division in Arabidopsis. *Plant Physiol.* **2012**, *160* (1), 332–348. https://doi.org/10.1104/pp.112.198705.
(22) Vogel, J. P.; Schuerman, P.; Woeste, K.; Brandstatter, I.; Kieber, J. J. Isolation and Characterization of Arabidopsis Mutants Defective in the Induction of Ethylene Biosynthesis by Cytokinin. *Genetics* **1998**, *149* (1), 417–427. https://doi.org/10.1093/genetics/149.1.417.

(23) Yang, L.; Xie, M.; Wu, Y.; Cui, X.; Tang, M.; Yang, L.; Xiang, Y.; Li, Y.; Bai, Z.; Huang, J.; Cheng, X.; Tong, C.; Liu, L.; Liu, S.; Zhao, C. Genetic Mapping and Regional Association Analysis Revealed a *CYTOKININ RESPONSE FACTOR 10* Gene Controlling Flowering Time in *Brassica Napus* L. *Ind. Crops Prod.* **2023**, *193*, 116239. https://doi.org/10.1016/j.indcrop.2023.116239.
(24) Li, X.; Fang, S.; Chen, W.; Liu, S.; Zhao, L.; Xu, Z.; Chen, S.; Liu, Y.; Du, Y.; Deng, L.; Liu, L.; Wang, T.; Li, P.; Zhu, Y.; Yu, D.; Wang, H. CRF12 Specifically Regulates the Flowering Time of Arabidopsis Thaliana under Non-Inductive Conditions. *Plant J.* **2025**, *121* (3), e17257. https://doi.org/10.1111/tpj.17257.
(25) Gaillochet, C.; Stiehl, T.; Wenzl, C.; Ripoll, J.-J.; Bailey-Steinitz, L. J.; Li, L.; Pfeiffer, A.; Miotk, A.; Hakenjos, J. P.; Forner, J.; Yanofsky, M. F.; Marciniak-Czochra, A.; Lohmann, J. U. Control of Plant Cell Fate Transitions by Transcriptional and Hormonal Signals. *eLife* **2017**, *6*, e30135. https://doi.org/10.7554/eLife.30135.
(26) Takei, K.; Sakakibara, H.; Sugiyama, T. Identification of Genes Encoding Adenylate Isopentenyltransferase, a Cytokinin Biosynthesis Enzyme, in*Arabidopsis Thaliana* *. *J. Biol. Chem.* **2001**, *276* (28), 26405–26410. https://doi.org/10.1074/jbc.M102130200.
(27) Kim, H. J.; Chiang, Y.-H.; Kieber, J. J.; Schaller, G. E. SCFKMD Controls Cytokinin Signaling by Regulating the Degradation of Type-B Response Regulators. *Proc. Natl. Acad. Sci.* **2013**, *110* (24), 10028–10033. https://doi.org/10.1073/pnas.1300403110.
(28) Hamant, O.; Nogué, F.; Belles-Boix, E.; Jublot, D.; Grandjean, O.; Traas, J.; Pautot, V. The KNAT2 Homeodomain Protein Interacts with Ethylene and Cytokinin Signaling. *Plant Physiol.* **2002**, *130* (2), 657–665. https://doi.org/10.1104/pp.004564.
(29) Ye, L.; Wang, X.; Lyu, M.; Siligato, R.; Eswaran, G.; Vainio, L.; Blomster, T.; Zhang, J.; Mähönen, A. P. Cytokinins Initiate Secondary Growth in the *Arabidopsis* Root through a Set of LBD Genes. *Curr. Biol.* **2021**, *31* (15), 3365-3373.e7. https://doi.org/10.1016/j.cub.2021.05.036.
(30) Makino, S.; Kiba, T.; Imamura, A.; Hanaki, N.; Nakamura, A.; Suzuki, T.; Taniguchi, M.; Ueguchi, C.; Sugiyama, T.; Mizuno, T. Genes Encoding Pseudo-Response Regulators: Insight into His-to-Asp Phosphorelay and Circadian Rhythm in Arabidopsis Thaliana. *Plant Cell Physiol.* **2000**, *41* (6), 791–803. https://doi.org/10.1093/pcp/41.6.791.
(31) Matsushika, A.; Makino, S.; Kojima, M.; Mizuno, T. Circadian Waves of Expression of the APRR1/TOC1 Family of Pseudo-Response Regulators in Arabidopsis Thaliana : Insight into the Plant Circadian Clock. *Plant Cell Physiol.* **2000**, *41* (9), 1002–1012. https://doi.org/10.1093/pcp/pcd043.
(32) Zhang, J.; Wrage, E. L.; Vankova, R.; Malbeck, J.; Neff, M. M. Over-Expression of SOB5 Suggests the Involvement of a Novel Plant Protein in Cytokinin-Mediated Development. *Plant J.* **2006**, *46* (5), 834–848. https://doi.org/10.1111/j.1365-313X.2006.02745.x.
(33) Sun, S.; Yu, J.-P.; Chen, F.; Zhao, T.-J.; Fang, X.-H.; Li, Y.-Q.; Sui, S.-F. TINY, a Dehydration-Responsive Element (DRE)-Binding Protein-like Transcription Factor Connecting the DRE- and Ethylene-Responsive Element-Mediated Signaling

Pathways in *Arabidopsis**. *J. Biol. Chem.* **2008**, *283* (10), 6261–6271. https://doi.org/10.1074/jbc.M706800200.
(34) Wang, J.; Ma, X.-M.; Kojima, M.; Sakakibara, H.; Hou, B.-K. N-Glucosyltransferase UGT76C2 Is Involved in Cytokinin Homeostasis and Cytokinin Response in Arabidopsis Thaliana. *Plant Cell Physiol.* **2011**, *52* (12), 2200–2213. https://doi.org/10.1093/pcp/pcr152.
(35) Li, Y.; Wang, B.; Dong, R.; Hou, B. *AtUGT76C2*, an *Arabidopsis* Cytokinin Glycosyltransferase Is Involved in Drought Stress Adaptation. *Plant Sci.* **2015**, *236*, 157–167. https://doi.org/10.1016/j.plantsci.2015.04.002.
(36) Gan, Y.; Liu, C.; Yu, H.; Broun, P. Integration of Cytokinin and Gibberellin Signalling by Arabidopsis Transcription Factors GIS, ZFP8 and GIS2 in the Regulation of Epidermal Cell Fate. *Development* **2007**, *134* (11), 2073–2081. https://doi.org/10.1242/dev.005017.
